# Germline restricted chromosomes in dark winged fungus gnats show dynamic chromosome organisation despite strong purifying selection on genes

**DOI:** 10.1101/2025.10.24.684428

**Authors:** Christina N. Hodson, Aleksandra Bliznina, Federico Abascal, Sam Ebdon, Thomas C. Mathers, Raquel Vionette Do Amaral, Andrew Pomiankowski, Kamil S. Jaron

## Abstract

Germline restricted chromosomes (GRCs) are an evolutionary mystery. While they seem to be important and perhaps indispensable in species they occur, they also show surprising variability in copy number and size. We are just beginning to understand the evolutionary significance of these chromosomes. One factor that impedes our understanding is that at present, few clades with these chromosomes have been studied in depth. We genomically investigate the GRCs in dark-winged fungus gnats (Sciaridae), a cosmopolitan and species-rich taxa which have harboured GRCs for at least 58 million years. Sciaridae species carry from 1-4 large GRCs which evolved through introgression from the Cecidomyiidae lineage. We produce the first male germline chromosome-scale genome assemblies for the sciarids *Bradysia coprophila, B. impatiens,* and *Lycoriella ingenua*. Comparing the GRCs both between and within species, we find that these chromosomes evolve in an extraordinarily dynamic fashion, with very little conservation of synteny or gene content. Puzzlingly, despite this variability, GRCs are gene rich and their genes are evolving under strong purifying selection. Additionally, we uncover evidence that somatic elimination of GRCs potentially occurs via a satellite mediated process. Our study sheds light on the mechanisms of meiosis, tissue restriction, chromosome elimination, and gives us a better appreciation for the diversity of genetic systems that not only persist, but thrive in nature.

**Significance statement:** Germline restricted chromosomes (GRCs) are a fascinating evolutionary puzzle. They are restricted to the germline early in development, have unusual transmission dynamics and are more variable than core chromosomes, but also seem to be indispensable in clades they occur in. We generate chromosome level GRC assemblies from three species of fungus gnats (Sciaridae, Diptera). We find that the GRCs, which originated via a cross-family hybridization event, are evolving much more dynamically than other chromosomes in the genome. The five GRCs we sequence are all large, gene-rich, and seem to carry functional genes, but contain mostly a non-overlapping gene set, suggesting that four of the five GRCs we sequenced did not evolve from the same original chromosome.

## Introduction

Some chromosomes are found exclusively in the germ cells, as they are programmed to be eliminated from somatic cells early in development (Gerbi, 2022; Wang and Davis, 2014). Known as germline restricted chromosomes (GRCs), they are a mystery from an evolutionary standpoint. They have evolved in at least six different lineages and have persisted in nearly all members of these large clades for more than 50 million years (Bauer and Beermann, 1952; Kahle, 1908; Metz et al., 1926; Nakai et al., 1991; Pigozzi and Solari, 1998). This suggests that they have a conserved indispensable function in these clades. However, contrary to these expectations, there is variability in both GRC number, size, and gene content between even closely related species (Hodson and Ross, 2021; Ruiz-Ruano et al., 2025; Schlebusch et al., 2023; Torgasheva et al., 2019). Therefore, there is debate whether GRCs provide an adaptive advantage (Smith et al., 2018), act selfishly (Pei et al., 2022) or potentially both (Vontzou et al., 2023). GRCs restriction to the germline occurs early in development in a highly programmed manner, generally involving the GRCs being left on the metaphase plate in early embryonic divisions and not being incorporated into the somatic cell lineage, but behaving typically in the germ cell lineage (de Saint Phalle and Sullivan, 1996; Geyer-Duszyńska, 1959; Staiber, 2006; Timoshevskiy et al., 2016). How this process occurs mechanistically, how it evolves, and how being restricted to the germline shapes chromosome evolution is largely unknown. Furthermore, GRCs show non-Mendelian inheritance in all of the well-studied clades (songbirds- Pigozzi and Solari, 2005; fungus gnats- Rieffel and Crouse, 1966; gall gnats- White, 1973). This is perhaps a way to regulate GRC number, but the evolution of these non-canonical transmission dynamics in the context of the other features mentioned is unclear. Therefore, the unusual combination of features of GRCs is challenging our views on how conserved functions are encoded genomically, and opens an opportunity to understand more general principles of genome evolution.

One factor impeding our understanding of GRC evolution, and how it fits into the wider context of chromosome evolution and meiosis, is the fact that we lack detailed information about GRCs from multiple clades. Flies with GRCs have been well-studied cytogenetically (Escribá et al., 2011; Gerbi, 2022; Metz, 1938; White, 1973), but until recently minimal genomic work has been undertaken on this group (Crane et al., 2023; Hodson et al., 2022). Interestingly, all three fly clades with GRCs (Sciaridae, Cecidomyiidae, and Chironomidae) are extremely species rich, noted as dark-taxa, where molecular evidence from DNA barcoding suggests that the number of species in the family is severely under-described (Meier et al., 2025). In all three families, GRCs are expected to have evolved through polyploidization. In Chironomidae and Cecidomyiidae, it is thought that GRCs, which are generally numerous within individuals (i.e. several times more abundant than the set of core chromosomes found in all cells) evolved through several rounds of whole genome duplication (White, 1973). In contrast, GRCs in Sciaridae were recently shown to have originated through a cross-family hybridization event with a Cecidomyiidae individual, with the GRCs being of Cecidomyiidae origin and the core chromosomes of Sciaridae origin (Hodson et al., 2022). GRCs in Sciaridae are much less numerous, ranging in number from 0-4 (note that GRC loss has only been recorded in one species), with most species carrying two GRCs (Gerbi, 1986; Hodson and Ross, 2021). This, combined with the observation that GRCs form a bivalent in female meiosis (Rieffel and Crouse 1966), led to the assumption that a single Cecidomyiidae chromosome introgressed into the Sciaridae lineage and became the GRCs. This hypothesis would suggest GRCs in Sciaridae are similar to Passarine birds, where the single GRC in different species share the same origin (Ruiz-Ruano et al., 2025). However, most research on Sciaridae GRCs, including the recent genomics paper (Hodson et al., 2022), focused on one species, *Bradysia coprophila*, and in this species there was coverage differences of GRC sequences between the two chromosomes, suggesting that not all regions of the GRCs are found on both chromosomes. Therefore, we need to sequence a broader array of Sciaridae species to confirm 1. Whether all GRCs across the family show evidence of introgression from Cecidomyiidae, 2. Whether GRCs in different species share a common origin and 3. Whether two GRCs within species are homologous.

In addition, GRCs in Sciaridae have some notable similarities to the sex chromosomes in this family. Sciaridae has an XO sex chromosome system, however, sex is not determined by the number of X chromosomes inherited from parents but by the number of X chromosomes eliminated in somatic cells in early embryogenesis, which is under maternal control (Du Bois, 1933; C. W. Metz, 1938). The elimination of X chromosomes in early embryogenesis looks very similar to GRC elimination, with the exception that it happens a few cell divisions later in embryonic development (de Saint Phalle and Sullivan, 1996). Therefore, these two elimination events are potentially related. The X chromosome elimination in embryogenesis is associated with the short arm of the acrocentric X chromosome (Crouse, 1977). An rDNA-rich region within the short arm, designated the controlling element, is known to drive the elimination process during embryogenesis, since translocating this region to another chromosome results in the elimination of that chromosome, rather than the X (Crouse, 1979, 1977). It was initially hypothesized that the GRCs evolved from the X chromosome because of the similarities in how they are eliminated and the fact that both have unusual transmission dynamics through males (Haig, 1993). It is now known that GRCs did not evolve from the X chromosome (Hodson et al., 2022), but whether the mechanism of somatic elimination could be related between the two chromosome types, and how, is an open question.

Finally, one large unanswered question regarding GRCs in Sciarids is whether these chromosomes have a germline function. In other clades with GRCs, these chromosomes have been shown to carry genes enriched in functions relating to reproduction (Kinsella et al., 2019; Vontzou et al., 2025; Yasmin et al., 2022). However, a major difference between the *B. coprophila* GRCs and the GRCs in Passarine birds and lampreys is that the GRCs in *B. coprophila* carry more than 10,000 genes rather than a few hundred, and so far conducting gene expression analyses for the GRCs has been challenging, showing little expression of genes on these chromosomes (Kyriacou et al., 2025). Detailed studies of chromosome appearance in *B. coprophila* over development suggests that the two GRCs in *B. coprophila* are heterochromatic at several stages of development (Rieffel and Crouse, 1966), leading to the speculation that these chromosomes may carry few important genes (Charles W. Metz, 1938). However, since GRCs have been retained in sciarids for over 50 million years, they likely have genes performing an important function in this lineage. Comparing GRC content between multiple species, as well as comparing sequence evolution between GRC genes and genes in the rest of the genome (core chromosomes) can help up understand whether GRC genes have an important function, or whether they are truly selfish chromosomes which managed to persist for tens of millions of years.

We generate the first chromosome level germline genomes from three Sciaridae species to delve into unresolved questions regarding their evolution. We sequence *B. coprophila*, *B. impatiens,* and *Lycoriella ingenua,* with two, one and two GRCs, respectively. We characterise the GRCs in these three species genomically and explore how similar GRCs are between species. We find that each of the five GRCs we sequence is largely distinct, with the exception of GRC1 in *B. coprophila* and GRC2 in *L. ingenua,* which share two large syntenic regions. All five GRCs are mosaics of regions originating from either Cecidomyiidae or Sciaridae, supporting a hybrid origin of GRCs with more recent duplications from core sciarid chromosomes onto GRCs. Interestingly, we find that 1. species with two GRCs carry two non-homologous chromosomes 2. genes on the GRCs are under strong purifying selection similar to core chromosome genes and 3. GRCs may share a similar elimination mechanism to the X chromosome in *B. coprophila*. Therefore, the GRCs in sciarids contain thousands of seemingly functional genes, but counterintuitively gene content is distinct between different GRCs in the same species and between species.This study provides the first genomic comparison of GRCs in multiple fly species. As such, it provides an important resource for future work. Understanding the nature of GRC evolution in flies and in other clades will enable us to place GRCs in a broader evolutionary framework. Given the mystery surrounding how GRCs evolve, this has the potential to shape our ideas about chromosome evolution in a substantial way.

## Results

### Characteristics of GRCs in fungus gnats

We generated chromosome level genomes for *B. coprophila*, *B. impatiens,* and *L. ingenua* using high-coverage PacBio HiFi and Hi-C data derived from the pooled testes of siblings from inbred lines maintained in the laboratory. To classify chromosomes as autosomes, X chromosomes, or GRCs, we used a previously described method that accounts for differences in *k*-mer coverage profiles between the core chromosomes and the GRCs, when comparing sequencing data derived from germline to GRC-less samples (somatic tissue or a whole organism) (Hodson et al., 2022). As a result, each of the genome assemblies contain four core chromosomes (three autosomes and one X chromosome), and either two GRCs in the case of *B. coprophila* and *L. ingenua*, or one GRC in case of *B. impatiens*. The karyotypes inferred from the assemblies were consistent with previous cytological determinations for both *Bradysia* species (Metz 1926; 1938; Carson 1944, **Figure 1**), except the two GRCs were suggested to be homologous chromosomes (Crouse et al., 1971), but in *B. coprophila* (with two GRCs), the two chromosomes showed little to no homology to each other. Furthermore, the two GRCs for *L. ingenua* (Metz, 1926; Metz et al., 1926) are described as large V-shaped chromosomes in the cytological literature (**Figure 1**), while we sequenced two chromosomes that are significantly smaller in size compared to the core chromosomes. That might indicate that the lab population in these studies differed from ours, suggesting population variability in GRCs within *L. ingenua,* or that large scale changes in the GRCs occurred in this species since these initial studies. As chromosome names are only available for *B. coprophila*, we decided to keep the naming of chromosomes by size for all three genomes for consistency in this paper (**Supplementary Table 1** for correspondence between *B coprophila* chromosome names in the literature and our assembly; Crouse, 1943; Urban et al., 2025).

**Figure 1.**
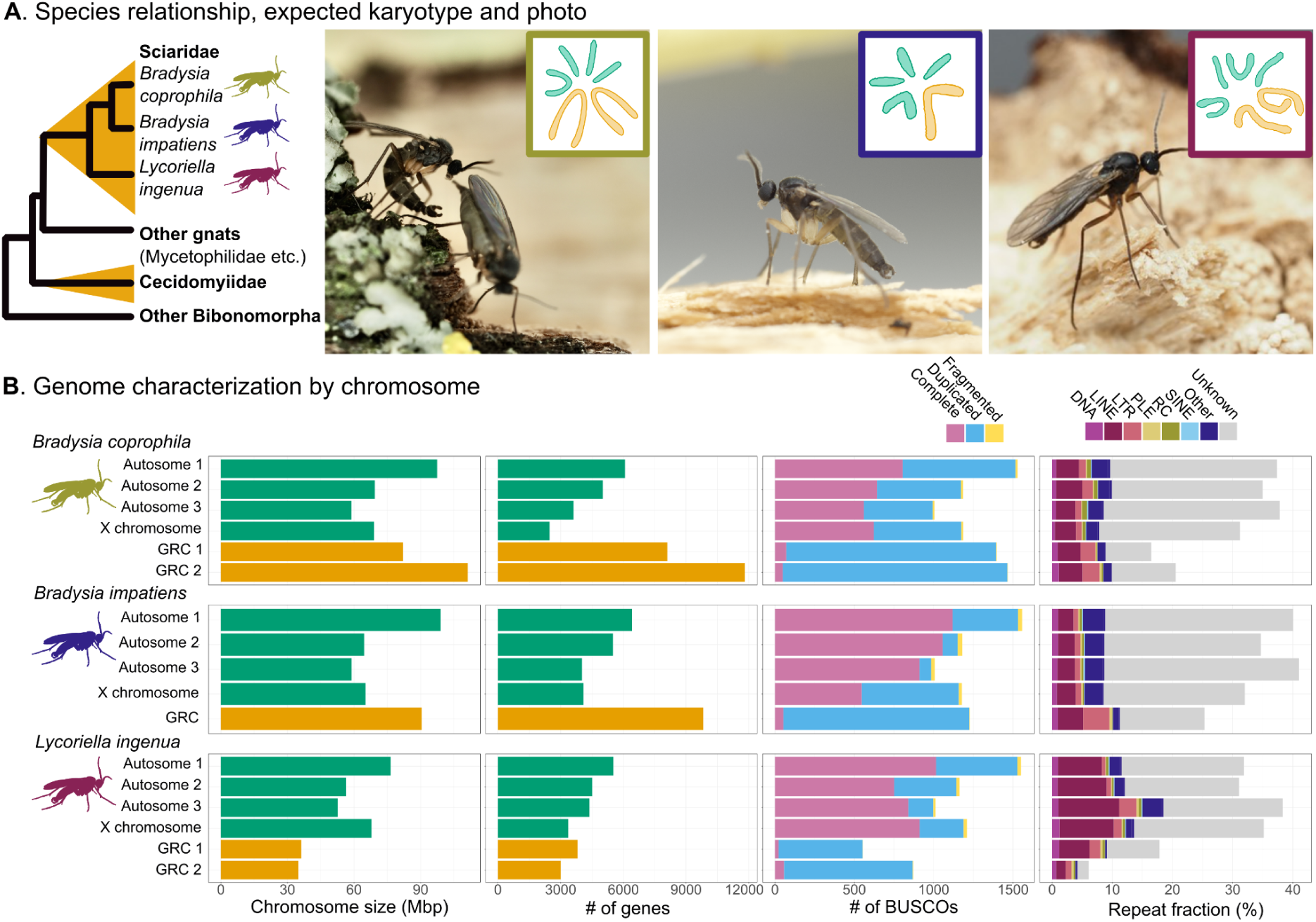
Chromosome constitution and genomic features in the three sciarid species. **A**. Cladogram showing the phylogenetic position of the focal species (the orange triangles indicate clades with GRCs) and images of *B. coprophila* (left)*, B. impatiens* (middle), and *L. ingenua* (right) with the haploid chromosome constitution inset (chromosome images adapted from Metz 1926; DuBois 1932; Carson 1944). The germline restricted chromosomes are coloured orange (large V shaped chromosomes in all species), while the core chromosomes (X chromosome and autosomes) are coloured green. Coloured borders on the chromosome inset will be used throughout the remainder of the paper to distinguish each species. **B**. Chromosome features for each chromosome within each species from our germline genome assemblies. Chromosome size, number of genes on each chromosome, number of conserved dipteran BUSCO genes (n=5067), and proportion of each chromosome made up of repeats of various classes are shown (note: The other class in the TE plot contains low-complexity sequences, satellites, and simple repeats). Photo credit: S. Ebdon/C. Hodson.

The core chromosomes show clear synteny between the three species (**Supplementary Figure 1**), and are consistent in size. The largest autosome of each species ranges in size from 76.5 Mbp in *L. ingenua* to 99.1 Mbp in *B. impatiens*, and the smallest chromosome ranges in size from 52.4 Mbp in *L. ingenua* to 58.9 Mbp in *B. impatiens* (**Figure 1**). Unlike the core chromosomes, the GRCs vary a lot in size: 35 and 36 Mbp in *L. ingenua*, 90 Mbp in *B. impatiens*, and 82 and 111 Mbp in *B. coprophila*. This variation explains the difference in the total genome size observed between the species (GRCs in *B.coprophila* correspond to around 40% of the genome, while in *L. ingenua* and *B.impatiens* it is 21-23%; **Figure 1B**). The reference genomes are highly complete with 94.8-97.1% of complete BUSCOs (Benchmarking Using Single Copy Orthologues) with the “diptera_odb12” dataset (**Supplementary Table 2**), with a large proportion of duplicated BUSCOs, as expected if the GRCs contain genes also present on the core chromosomes (**Figure 1B**). Moreover, each GRC has thousands of annotated protein-coding genes, ranging between 3,006 in *L. ingenua* GRC2 to 11,816 in *B. coprophila* GRC2.

Repeat annotation of the genomes showed depleted TE content in the GRCs compared to the core chromosomes, ranging from 6.8% for *L. ingenua* GRC2 to 25.3% for *B. impatiens* GRC (vs. >30% TEs for all core chromosomes, **Figure 1B**). GRCs in all genomes are enriched in LINE, LTR and DNA transposable elements (% chromosome in core vs. GRC: LINE: 4.53 vs. 4.70; LTR: 1.04 vs. 3.46 DNA: 0.80 vs. 1.22). Among them, TEs from LTR and DNA subclasses are enriched in and shared between all GRCs. The sequence divergence of some TE groups in the GRCs, such as unknown and DNA transposons, has a bimodal distribution indicating presence of both ancient and recently active (or recently acquired) transposons, especially in *B. impatiens* (**Supplementary Figure 2**).

### All GRCs originated via introgression from Cecidomyiidae and show high gene turnover

Previous work (Hodson et al., 2022) suggested that the GRCs in *B. coprophila* arose through a hybridization event between an ancient sciarid and cecidomyiid leading to germ-line restricted allopolyploid. We revisited this hypothesis with our three chromosome level genomes, and analysed the evolution of the germ-line restricted subgenome. We analysed between 436-1077 BUSCOs for each of the GRC chromosomes. In agreement with Hodson 2022, we identified the same two dominant topologies for all GRCs - hybrid origin genes (placed in Cecidomyiidae clade) and recent duplications from the core chromosomes (placed within Sciaridae clade) (**Figure 2B**). Very few genes had other placements (6 total; note that many genes have uncertain placements due to either the Cecidomyiidae or Sciaridae family not being monophyletic in the gene tree). In contrast, the core genes of the sequenced species nearly always fell within the Sciaridae clade in phylogenies (4661 out of 4666 analysable gene trees). GRC gene placement in phylogenies supports the hypothesis that GRCs within Sciaridae evolved through an ancient hybridization between a cecidomyiid and sciarid, with the cecidomyiid contributing the GRCs, followed by duplications of blocks of sequence from the core chromosomes (of sciarid origin) onto the GRC (See **Supplementary Figure 4** for hybridization scheme). Assuming a single origin of GRCs, we expected the introgressed BUSCO genes to have the same ancestry and form a monophyletic subclade within the Cecidomyiidae family. However, we observed that the five GRCs we sequenced show three distinct placements in the phylogeny (**Figure 2D**). The topology of individual gene-trees revealed that all GRCs show similar patterns in terms of the most frequent nearest non-GRC neighbour (**Figure 2D** - inset). For all GRCs, the most common non-GRC neighbour was the cecidomyiid aphid killer *Aphidoletes aphidimyza*, although for the GRC in *B. impatiens* this was not the closest non-GRC neighbour in the species phylogeny (**Figure 2D)**. When compared to the aphid killer *A. aphidimyza* genome, different GRCs have higher similarity to different core chromosomes in that genome, suggesting a more complex hybridisation scenario. For example, the GRCs we sequenced could have evolved from different core chromosomes in Cecidomyiidae (**Supplementary Figure 5).** Alternatively, placement of individual GRC branches in the phylogeny could be due to technical artefact resulting from the differences in gene content between the GRCs (Roure et al., 2012).

**Figure 2.**
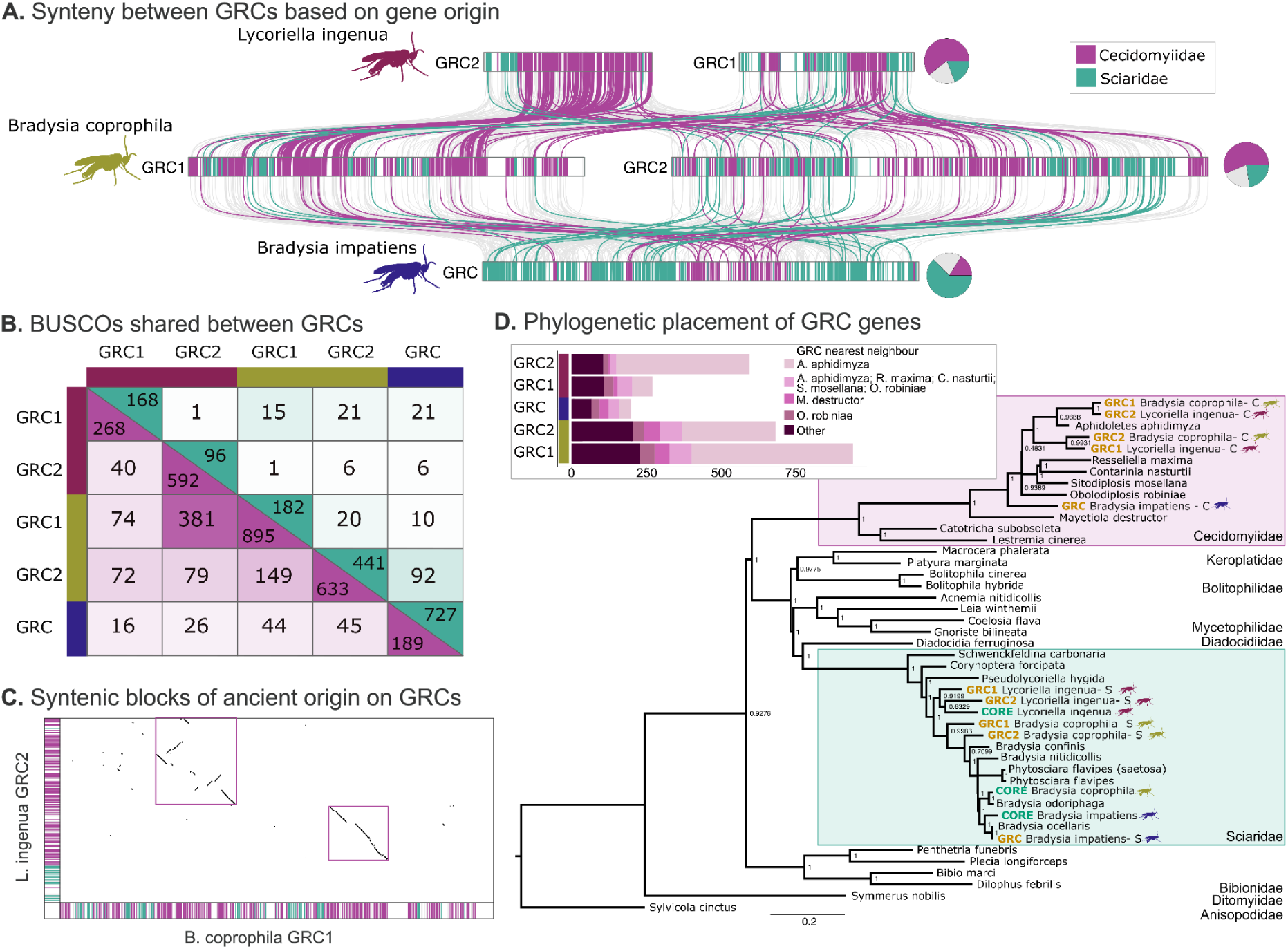
Five distinct germline restricted chromosomes in *B. coprophila, B. impatiens and L. ingenua*. **A.** Synteny ribbon plot for GRCs in *L. ingenua* (top-pink), *B. coprophila* (middle-yellow) and *B. impatiens* (bottom-blue) (See **Supplementary** Figure 1 for the corresponding plot for core chromosomes). The origin of BUSCO genes according to phylogenetic placement is shown along each chromosome (Sciaridae origin genes = teal, Cecidomyiidae origin genes = purple). Lines are connecting the same BUSCO on different GRCs, with lines coloured if the genes share the same origin (and therefore potentially related by descent) in each species. The pie chart indicates the proportion of BUSCOs with a Cecidomyiidae, Sciaridae or other (mostly unknown) origin for each species. **B.** Number of BUSCOs with the same origin between two GRCs. Sciaridae origin genes are shown on the top (teal) and Cecidomyiidae origin genes are shown on the bottom (purple). The numbers on the diagonals are the number of BUSCOs of each origin on each chromosome in total. *L. ingenua* GRCs are indicated with pink, *B. coprophila* GRCs are indicated in yellow, *B. impatiens* with blue. **C.** Synteny between GRC1 in *B. coprophila* and GRC2 in *L. ingenua*. These two chromosomes share the largest number of BUSCO genes of cecidomyiid origin and show clear synteny between the two chromosomes, with two syntenic regions of approximately 10 (left) and 7.3 (right) Mbp. **D.** Species tree showing the phylogenetic position of each GRC partitioned by the phylogenetic origin of BUSCOs. Numbers at the nodes indicate the posterior probability at that node. The inset shows a barplot of the nearest non-GRC neighbour in gene-trees for each gene with a cecidomyiid origin. Colours on the left of each bar indicate species identity and match colouring from **A.**

Our initial expectation was clear homology and similarity of GRCs subgenomes of the three relatively closely related species. However, GRCs share relatively few genes of Cecidomyiid origin in common (**Figure 2B**), such that for the vast majority of GRC BUSCOs, there would be a copy on some GRCs but missing data for others. We found that there was very little BUSCO synteny between GRCs, in sharp contrast to the clear synteny between core chromosomes of the three species (**Supplementary Figure 1**). An average of 92 BUSCOs of Cecidomyiidae origin were shared between any two GRCs (and therefore potentially shared through descent). GRC1 of *B. coprophila* and GRC2 of *L. ingenua* shared 381 BUSCOs of Cecidomyiidae origin (**Figure 2B** and **C**). These two chromosomes have two regions of clear synteny approximately 10 Mbp and 7.3 Mbp respectively, which contain BUSCOs of cecidomyiid origin, suggesting that these chromosomes have the same origin and retain genomic signatures of that origin (**Figure 2A** and **C**).

The genes on the GRCs are arranged in blocks of shared origin along the chromosomes (**Figure 2A**), and the number and arrangement of blocks varies between GRCs. Most sciarid-origin genes form a clade closely related to the core gene of the same species (**Figure 2D**), showing the dynamic nature of evolution on the GRCs (**Supplementary Figure 6**). Notably, the *B. impatiens* GRC has a substantially larger number of sciarid origin BUSCOs and sciarid origin blocks along the GRC (**Figure 2A**) compared to other GRCs. On the other hand, core genomes showed only very limited segmental duplications of the genes of hybrid origin **(Supplementary Figure 3)**. There were five cases where a core Sciaridae gene from one of the three focal species was found in the Cecidomyiidae clade (0.12% of 4139 BUSCO gene trees), raising a question about mechanism preventing gene movement in the opposite direction. We hypothesise that the turnovers via segmental duplications from the core genome on to GRCs are an ongoing process that resulted in differential loss of the hybrid origin genes in different lineages.

### GRC genes do not show evidence of sequence degradation

In absence of clear overlapping hybrid gene sets, we hypothesised the gene content of the GRCs will be subjected to relaxed purifying selection and show signatures of decay. To test this hypothesis, we identified orthologous groups in the genome to compare genes on the GRCs to homologs in the core genome, as well as *Pseudolycoriella hygida*, *Obolodiplosis robiniae, A. aphidimyza, Contarinia nasturtii*, and *D. melanogaster.* We examined evolutionary evidence of GRC gene functionality (using genomic metrics) over time. We did this in a few different ways depending on the type of gene. We compared pairs of homologous genes on different chromosomes given origin using dN/dS analysis, and more detailed examination of large recent segmental duplications between core and GRCs exploiting persisting synteny to analyse number of exons and lengths of genes for signatures of truncating mutations on the GRCs.

We stratified GRC genes by their evolutionary origin and estimated the strength of purifying selection acting on them for 5674 orthogroups containing at least one homologous core and GRC gene pair and for which we could identify the origin of the GRC gene (i.e. Cecidomyiidae or Sciaridae). We analysed whether the dN/dS in the equivalent core gene branch was different from the GRC gene branch depending on the gene origin (see methods/ **Supplementary figure 7** to schematic of what an equivalent branch meant in the case that the gene was of cecidomyiid vs. sciarid origin). Our results showed that GRC genes are evolving under very strong purifying selection (mean dN/dS of 0.054 for core genes of cecidomyiid origin vs 0.058 for GRC genes, and 0.050 vs. 0.066 for sciarid origin core and GRC genes, respectively), and comparable, although slightly higher, than core genes. We found no difference in the dN/dS between GRC and core gene branches for GRC genes in the Cecidomyiidae clade (paired t-test: t = -1.67, df= 1614, p= 0.09, **Figure 3A**), but the GRC gene branches had a higher dN/dS than the equivalent core branches for genes in the Sciaridae clade (paired t-test: t = -5.42, df= 1275, p= 7.2e-8, **Figure 3A**). We also compared the dN/dS for each GRC terminal branch compared to the entire gene tree for each orthogroup, and found that GRC branches had similar dN/dS values as the entire gene tree, both for genes of Cecidomyiidae and Sciaridae origin (**Supplementary Figure 8**).

**Figure 3.**
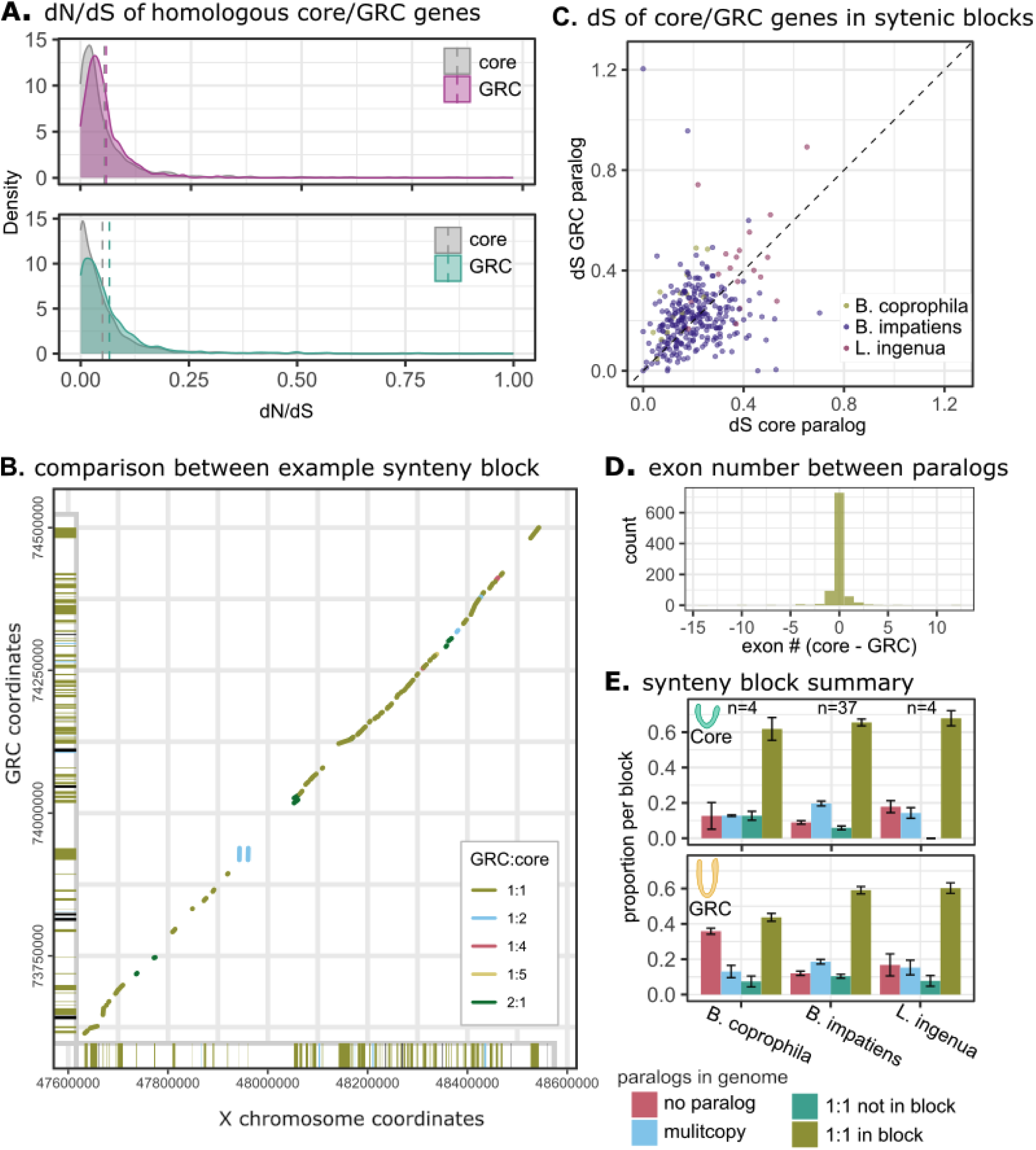
Comparison of homologous genes on the GRCs to their core genome counterpart. **A**. Density plot of dN/dS values for GRC branches and the equivalent core gene branches for genes of Cecidomyiidae origin (top purple vs. grey for core) and Sciaridae origin (bottom teal vs grey for core). The means for each category are shown as dotted lines. **B**. An example synteny block between the GRC in *B. impatiens* and the X chromosome. This block contains 52 paralogs with one copy on the GRC and one in the core genome portion of the synteny block, out of 79 core and 79 GRC genes, respectively. The bars on the X and Y axis indicate whether each gene has a single paralog on the corresponding block (yellow), multiple paralogs (blue) or no paralog (black). In cases where there are multiple paralogs within the block, the genes in the main plot are coloured by the number of copies. **C**. dS values for the core gene (x-axis) and GRC gene (y-axis) with one copy on the GRC and one copy in the core genome in syntenic blocks of recent (sciarid) origin. Blocks in each species are indicated with different colours. **D**. Comparison of exon number between paralog pairs in syntenic blocks, with values at 0 indicating the same number of exons on the GRC copy of the gene as on the core genome copy. **E**. Summary of the proportion of genes in all synteny blocks analysed in each species. Values for the core genome counterpart of the synteny block are shown above the GRC portion. Genes with a copy on the focal counterpart (core-top or GRC-bottom) of the synteny block but not on the other chromosome are shown in pink, genes with multiple copies throughout the genome are shown in blue, genes with one paralog in the focal synteny block and exactly one paralog somewhere else in the genome but not on the core/GRC block (green), or genes forming a paralog pair, with one copy on the GRC counterpart of the synteny block and one copy on the core genome half of the synteny block (yellow- and no other homologs in the genome).

We identified large syntenic blocks between the core genome and the GRCs of Sciarid origin, which represent more recent duplication onto the GRCs from the core genome. These blocks allow us to do a more in depth comparison of gene features and gene presence between GRC regions of known ancestry and composition. We identified 37 blocks longer than 200 kbp and with more than 10 paralog pairs in *B. impatiens,* and 4 blocks in each of *B. coprophila* and *L. ingenua* (information about the blocks are in **Supplementary Table 3**). Given that the *B. impatiens* GRC carries a greater proportion of sequences of Sciarid origin than GRCs of other species we sequenced, it is not surprising that we found more synteny between the GRC and core genome in *B. impatiens*. We found that the GRC synteny blocks look very similar to their core counterparts, with most genes in the core genome block having one paralog on the GRC block, and no evidence that there is a surplus of missing GRC paralogs (i.e. genes on the core counterpart of the synteny block with no copy on the GRC synteny block) which would potentially represent degradation of GRC genes to the point that they are not annotated (**Figure 3B/E; Supplementary Table 4**). Although note that there are an excess of missing genes on the GRC portion of the synteny block. We believe this is due to poorer annotation of GRC genes due to lack of RNAseq data; for instance, TEs annotated as genes (**Supplementary Table 4**). Additionally, comparing the gene features of genes with one copy in the core synteny block and one copy on the GRC block shows that GRC genes have a similar exon number (paired t-test: t = -0.5653, df = 933, p-value = 0.572) and length (paired t-test: t = 1.6845, df = 933, p-value = 0.09242) to their core counterpart in the vast majority of cases, suggesting that compared to their core genome counterparts, these genes appear functional (**Figure 3D**; **Supplementary Figure 9**).

Despite low gene conservation of GRC chromosomes, our results provide indirect evidence of GRC genes being functional. Furthermore, we do not observe any form of gene decay in both old and recently duplicated genes. The dS values for genes found in syntenic blocks on GRCs and the core genome ranged from 0-1.2 (**Figure 3C**), which is similar to the range in which other studies have found gene degradation/pseudogenization on sex chromosomes (Akagi et al., 2025; Moraga et al., 2025) suggesting it is not due to limited time for gene decay to manifest. Finally, these results indicate that GRCs recombine, given that recombination is generally considered a fundamental prerequisite for efficient purifying selection in species with finite effective population sizes (Otto, 2021).

### GRC satellite evolution and its relationship to GRC recombination and somatic elimination

We characterised the satellite landscape in the genomes of *B. coprophila, B. impatiens,* and *L. ingenua,* to: 1) determine the location of the centromeres to analyse the centromere effect on recombination on core chromosomes and GRCs, 2) explore whether there is genomic evidence for GRCs pairing in female meiosis, and 3) examine the similarity between the controlling element on the X chromosome (contained in the short arm of the X chromosome near the centromere) and the GRCs.

We inferred the positions of candidate centromeres using GC-content and satellite abundance across individual chromosomes, and Hi-C off-diagonal signal between chromosomes (Methods). This approach identified a single putative centromeric locus on the autosomes and the X chromosome but not on the GRCs (**Supplementary Figure 10**). For GRCs, we chose the single best candidate centromeric locus (see methods for more detailed information). Using *k*-mer–based Jaccard similarity analyses, we confirmed that the centromeres of GRCs in *B. coprophila* and *L. ingenua* have distinct satellite compositions compared to the centromeres of the core chromosomes of the same species, consistent with previous cytological studies for *B. coprophila* (Kerrebrock et al. 2025; **Figure 4A**). In *B. coprophila*, both GRCs are metacentric and enriched with 156 bp and 185 bp satellite DNA, whereas the GRCs in *L. ingenua* contain a 126 bp satellite DNA near the chromosome ends and appear acrocentric, in contrast to earlier cytological reports (Metz, 1926). We further confirmed that neither BcopSat-156, BcopSat-185, nor LingSat-126 is present in the core chromosomes of their respective species, with the exception of *B. coprophila*’s X chromosome, which has ∼7 copies of BcopSat-185 on its short arm, covering approximately 1.3 kbp, the region near the putative location of the controlling element responsible for DNA elimination (**Figure 4B**; Crouse et al. 1977). Satellite analysis in *B. impatiens* revealed that centromeres on the autosomes and X chromosome share a 156 bp satellite DNA. Surprisingly, we also found this repeat at the end of the GRC near BUSCO genes with Sciaridae origin. Based on previous cytological studies in *B. impatiens* (Carson, 1944), we expected the GRC to be metacentric. We do observe a large array of GRC-specific 222/234 bp satellite DNA monomers in the middle of the *B. impatiens* GRC (**Supplementary Figure 11**). However, GC content patterns suggest the centromere is likely at the end of the chromosome (**Supplementary Figure 10**). Therefore, we believe that the GRC in *B. impatiens* was originally metacentric and composed of GRC exclusive satellites, but a new centromere might have been brought onto the GRC through a recent duplication from a core chromosome, suggesting a recent centromere turnover (**Supplementary Figure 12**).

**Figure 4.**
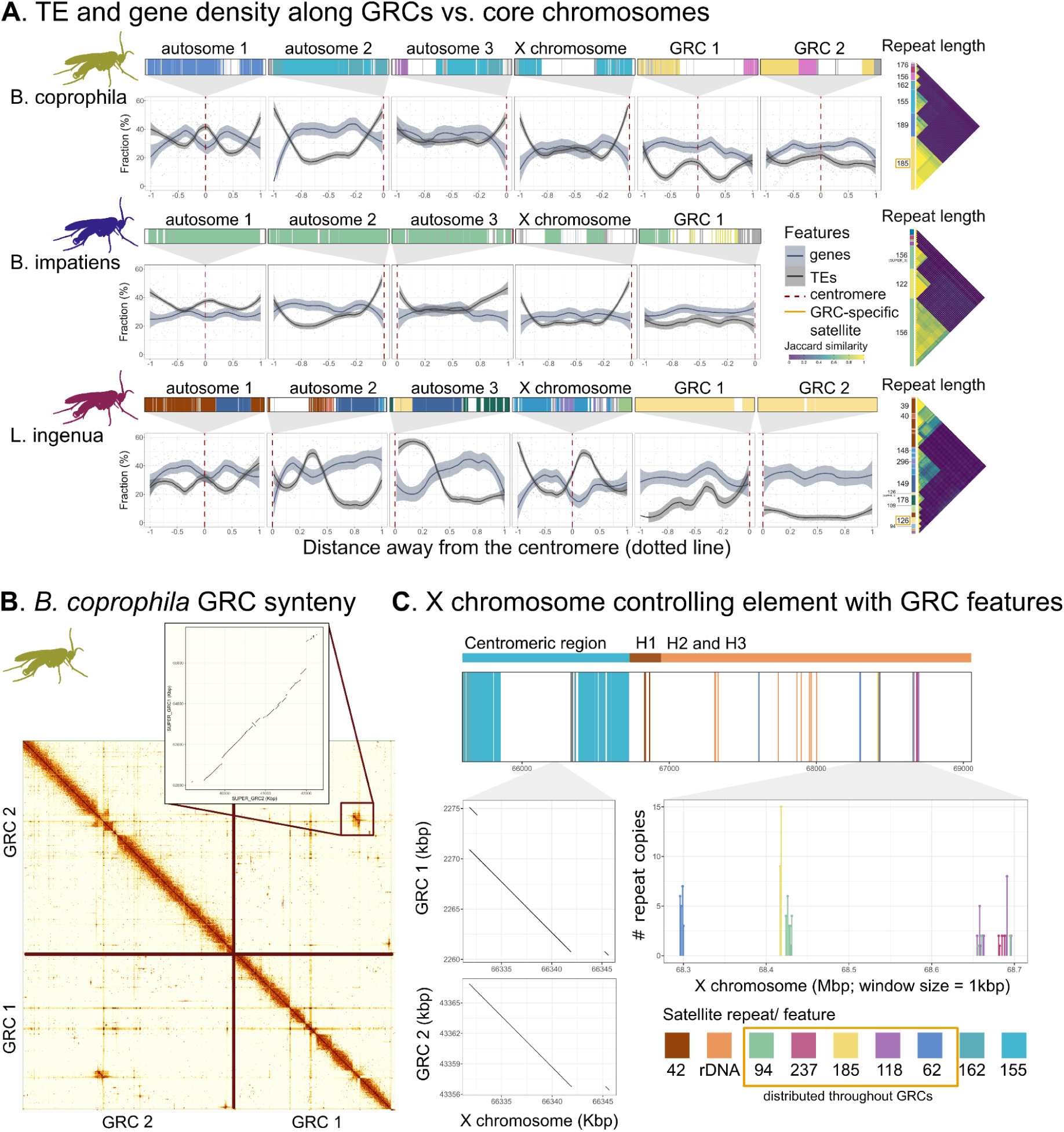
GRC homology and evolution through an analysis of satellite DNA. **A.** GRC centromere annotation and satellite composition for each species (top) and gene and TE density along each chromosome (bottom). Jaccard similarity between the repeats in the centromeric region is shown on the side of the plot, with GRC specific satellites indicated with an orange box. The satellites (both through identity and similarity) are different between the GRCs and core chromosomes in *B. coprophila* and *L. ingenua,* but the centromere in *B. impatiens* is made up of the core centromere satellite 156. Gene and TE density patterns also differ between the GRCs and core chromosomes, with most core chromosomes showing a pattern we would expect if recombination occurs (i.e. higher TE density and lower gene density near the centromere compared to the arm of the chromosome). **B.** Hi-C contact map for GRC1 and GRC2 in *B. coprophila*. The two GRCs have very little contact along most of their length, with an approximately 2 Mbp homologous region indicated through the off-diagonal signal (synteny shown in inset as a dotplot). **C.** Feature composition in the controlling element region of the X chromosome in *B. coprophila*. The top plot shows the short arm of the X chromosome, which contains the controlling element which governs X chromosome elimination in early embryogenesis (expected to be located in region H3; Crouse, 1977). Satellites within this region are otherwise GRC specific satellites, located throughout the two GRCs. There are also several small synteny blocks between the GRCs and X chromosome within this region. Sequence for the 42 bp repeat and rDNA sequence from Escriba et al., 2011 and Burke et al. 1993, respectively.

Recombination is typically suppressed in regions near to centomeres (Beadle, 1932). This ‘centromere effect’ leads to a spatial pattern of reduced gene density and enrichment of TEs near centromeres (Naish et al., 2021). We examined these two features with the expectation that in recombining chromosomes, TE density would increase and gene density would decrease near the centromere and telomeres compared to the chromosome arms. We found that this was the case for the core chromosomes in *B. coprophila* and *B. impatiens*, and for some of the core chromosomes in *L. ingenua* (**Figure 4A**). In *L. ingenua*, two of the core chromosomes (including the X chromosome) have unusual genomic feature distributions, which may be caused by recent structural rearrangements. In GRCs, TE and gene density were most often distributed more uniformly than in the core chromosomes. This may suggest that GRCs undergo more frequent structural rearrangements, which obscure expected patterns, or alternatively, the recombination landscape of GRCs does not follow typical patterns found on core chromosomes.

For both species with two GRCs, the two chromosomes share very few genes and limited synteny. We found only a small ∼2.1 Mbp region of homology between GRC1 and GRC2 of *B. coprophila* with 99.9% sequence identity (2.13 kbp of gaps; **Fig 3B**). Given these GRCs form a chromosome pair in female meiosis (Crouse et al., 1971), we speculate two possibilities. Either this 2.1 Mbp long homologous region between GRC1 and GRC2 in *B. coprophila* acts as a pairing region between these two chromosomes, or the two GRCs do not recombine (in which case if GRCs recombine in females than they would not carry both GRCs at the same time). The putative pairing region is depleted in gene content, but enriched in transposable elements and tandem repeats (**Supplementary Figure 13**). One 57 bp long satellite is found only in the homologous regions of the GRCs, and is a combination of satellite DNAs of 30 and 27 bp following each other. Moreover, the region is flanked by 94 bp repeat specific only to the GRCs and the short arm of the X chromosome (**Figure 4C**). Interestingly, the syntenic block is not located in the same region as the centromere in either chromosome. While this region presents an explanation for past observation of chromosomal pairing in two distinct, largely non-homologous chromosomes (the first scenario- Crouse et al., 1971), such explanation is puzzling in the light of strong purifying selection (that requires some form of recombination)(Otto, 2021). However, we also do not observe the centromere effect suggesting the recombination landscape might be unusual. Finally, the two scenarios are not mutually exclusive if females can carry either two of the same or two distinct GRCs.

Previous studies identified a controlling element on the short arm of the X chromosome in *B. coprophila* that is responsible for X-chromosome elimination (associated with sex determination) in early embryogenesis (Crouse et al., 1977). We wanted to see whether this region shared similarity to the two GRCs in this species, as GRCs are also eliminated in early embryogenesis in a manner similar to the X chromosome (de Saint Phalle and Sullivan, 1996). To do this, we searched for shared features between the X short arm and the GRCs. In addition to the mostly GRC-specific BcopSat-185 satellite DNA present in the short arm of the X chromosome, we identified four satellite DNAs of lengths 62, 94, 118, and 237 bp that are unique to and shared between the X short arm and the GRCs (**Figure 4C**). These satellites occur in low copy numbers and are localized near the end of the X chromosome, whereas in the GRCs they are distributed along the entire chromosome (**Supplementary Figure 14**). Furthermore, alignment of the X chromosome short arm with the GRCs revealed one synteny block (∼10 kbp) found in all three chromosomes (X, GRC 1, and GRC2). BLAST searches of these satellite repeats and syntenic regions against the rest of the genome confirmed their absence from the autosomes and the long arm of the X chromosome in *B. coprophila*. This suggests a potentially satellite mediated elimination mechanism for both the GRCs and X chromosome in early embryogenesis.

## Discussion

Using chromosome resolved germline genomes from three Sciaridae species, we have uncovered additional support for the finding that the germline restricted chromosomes in Sciaridae likely evolved through hybridization between an ancient cecidomyiid and the ancestor of all sciarids, with the GRCs arising from the chromosomes donated from the cecidomyiid ancestor (Hodson et al., 2022). This is a fascinating example of cross-family introgression, the scale of which has been rarely seen in nature. We would estimate that Sciaridae and Cecidomyiidae were already distinct families which had diverged for tens of millions of years when this hybridization event happened (Hodson et al., 2022). How exactly this hybridization event occurred and resulted in viable offspring is unclear. It resulted in a diploid chromosome set of sciarid chromosomes (making up the core chromosomes) and likely haploid set of cecidomyiid chromosomes (resulting in GRCs) (See **Supplementary Figure 4** for our speculative hybridization scenario). Interestingly, an example of hybridization between extraordinarily large divergences was also recently shown in another lineage with germline restricted DNA: the ciliate *Paramecium sonneborni* (Bénitière et al., 2026) (although note that ciliates have a macro and micronucleus and therefore not a true germline). It will be interesting to see if hybridization over large genetic distances is often associated with restriction of the introgressed genes to the germline, overcoming physiological constraints for compatibility.

The improved Sciarid GRC assemblies we generated from three species, has shed light into the dynamics of GRC evolution in a way which previous studies were unable to do. One major finding from our analyses is that the five GRCs we sequenced are largely unique in terms of gene content and structure. Only two GRCs show any large-scale homology of ancient origin. This is suggestive that the entire Cecidomyiid germline genome was integrated into Sciaridae, as we see synteny between the GRCs and all core chromosomes in *A. aphidimyza* (**Supplementary Figure 5**) and find many different BUSCO genes of Cecidomyiidae origin on the GRCs (**Figure 2**). However, cecidomyiids have four chromosomes in their core genome but also more than 10 germline restricted chromosomes in germ cells (e.g. *A. aphidimyza* has 10-23 GRCs; Gruzova and Batalova, 1993). Therefore, the details of which chromosomes were integrated into the Sciaridae lineage and retained is still unclear. It is possible that if the full cecidomyiid germline genome introgressed into the sciarid lineage, different chromosomes became fixed in different sciarid lineages. If this is the case, it seems more likely that the chromosomes retained in this interfamily hybridization would be the cecidomyiid GRCs, as these would have already possessed a mechanism to restrict themselves to the germline, avoiding some of the negative consequences of newly formed polyploid systems (Baduel et al., 2018). In any case, at most 4 GRCs were observed in any sciarid, begging the question, what happened to the other chromosomes if the entire cecidomyiid genome with GRCs introgressed into the sciarid lineage? Additionally, answering the question of whether the GRCs in Sciaridae are homologous to GRCs or core chromosomes in Cecidomyiidae is challenging, as cytological studies focusing on sequencing GRCs in cecidomyiids have so far come to the conclusion that the GRCs in this lineage are near-copies of the core chromosomes (Crane et al., 2023). High quality sequencing of Cecidomyiidae GRCs along with more extensive and more taxonomically diverse sequencing of Sciaridae GRCs, would go a long way to establishing the nature of the hybridization event that resulted in GRCs in Sciaridae.

We now have a better idea of how GRCs in Sciaridae evolve over time. For all five GRCs we see a backbone of genes of cecidomyiid origin (which we believe represents the original genes on these chromosomes) interspersed with blocks of genes of sciarid origin. Therefore, it appears over time segmental duplications from the core chromosomes accumulate on the GRCs as many of them show long syntenic blocks with conserved gene order and high sequence similarity with the core chromosomes (**Figure 3**). This pattern seems to be mostly unidirectional, as the core chromosomes carry very few genes of cecidomyiid origin (**Supplementary Figure 3**). The mechanism of duplication onto the GRCs from core chromosomes is still a matter for debate. It seems that this process must happen infrequently, since within our phylogenetic analyses, genes of sciarid origin were not often shared between GRCs, evolved at different times in the history of each species we sequenced, and from different core chromosomes for each GRC (**Fig 2B**; **Supplementary Figure 6; Supplementary Table 3**). However, whether this could happen through infrequent recombination events between the GRCs and core chromosomes or through some other mechanism is unclear.

Both males and females of *B. coprophila* carry two GRCs (reviewed in Gerbi 2022). Males pass both GRCs through sperm with no recombination, in females the GRCs were observed to form a bivalent (Crouse et al. 1971), and therefore could potentially recombine. Our analyses showed the two GRCs in males of *B. coprophila* and *L. ingenua* show very little homology to each other, and therefore GRC1 and GRC2 within a species are unlikely to recombine with each other. However, since we sequenced sperm from males, it is possible females do not carry both GRC1 and GRC2, but two copies of one of the two chromosomes, which could reconcile the finding that we have two divergent GRCs with the observation by Crouse et al. 1971. We did not observe molecular signatures of recombination caused by the centromere effect (Beadle, 1932; Naish et al., 2021): gene density was not clearly reduced and TE density nor increased near the centromeres on each analysed GRCs. However, it is possible that the dynamic evolution on the GRCs, where there appears to be frequent large-scale genomic changes, or the fact that TE load is reduced on the GRCs make such metrics difficult to interpret for GRCs. We also observed high gene density, low levels of TEs on GRCs compared to the recombining core chromosomes, and similar dN/dS values for GRC vs. core genes, which is the opposite of what would be expected of non-recombining chromosomes such as Y chromosomes (Charlesworth et al., 2000). We therefore need additional genomic or cytological data to clarify the nature of recombination in GRCs, but the fact that GRC genes are under strong purifying selection would suggest at least occasional recombination.

One fascinating aspect of GRC evolution, first identified by Kerrebrock et al. (2025), and confirmed in our analyses, is that the GRC satellites are largely different from the core chromosomes. We found this is the case even for the centromeric regions in *B. coprophila* and *L. ingenua*. In contrast, the GRC of *B. impatiens* appears to have recently acquired the same centromere satellite as the core chromosomes. In addition, the short arm of the X chromosome in *B. coprophila* contains satellites otherwise only found in the GRCs. Intriguingly, this is the region of the X chromosome known to control X chromosome elimination in early embryogenesis (Crouse et al., 1977; Escribá et al., 2011). It is tempting to speculate that the satellite composition is involved in the mechanism of somatic elimination, as X chromosome elimination in somatic cells (which is part of the process of sex determination in scairids) happens in a remarkably similar manner as GRC elimination from somatic cells (reviewed in Gerbi 2022). However, more work is needed on this topic encompassing a larger variety of techniques to establish how chromosome elimination occurs in Sciaridae. One interesting aspect of this finding from an evolutionary standpoint, however, is it was unclear whether GRC elimination evolved before or after X chromosome elimination for sex determination in Sciaridae. This result hints that GRC elimination evolved first.

An emerging pattern in multiple clades with GRCs is that these chromosomes carry functional genes which often play a role in germ cell development/ reproduction, as would fit their status as germline restricted genes. This has been found in birds (Kinsella et al., 2019; Vontzou et al., 2025) as well as in lampreys (Timoshevskiy et al., 2025; Yasmin et al., 2022). GRC expression is difficult to assess in Sciarids, as getting reliable germline expression data from such tiny flies is technically challenging (Kyriacou et al., 2025). We look at GRC gene evolution as a proxy for whether these genes are likely functional, finding that they have similar dN/dS to core genome homologs and suggesting that GRC genes may retain functional capacity. In sciarid flies, however, unlike these other clades which seem to have several hundred genes on the GRCs at most, we are able to annotate thousands of protein coding genes on the GRCs, even for L. ingenua, which carries the smallest GRCs of the three species we sequenced. Additionally, as the gene content between GRCs in different species is quite different, whether GRC genes are expressed and have germline specific roles is still somewhat of a mystery. Both sequencing more sciarids to see if there is a core set of retained GRC genes across species, as well as doing more in depth expression analyses, would help answer this question. In any case, in this respect GRCs appear to evolve in a different manner to selfish chromosomes such as B chromosomes, which generally carry few genes and retain mostly genes specific for maintaining transmission.

One major difference between lineages with GRCs is how they initially evolve. In birds, it’s thought that the single GRC found in all species investigated likely evolved from a B chromosome (Johnson Pokorná and Reifová, 2021; Kinsella et al., 2019). In contrast, in flies, in which there are several independent origins of GRCs, all have evolved through some form of polyploidy. In both Chironomidae and Cecidomyiidae, whole genome duplication likely gave rise to GRCs, while in Sciarids, we now believe that an allopolyploidization event gave rise to GRCs. However, even though the origins of GRCs differ between lineages, genomic comparisons have uncovered some similarities. Passerine birds, like Sciarids, have extraordinarily dynamic GRCs, which vary in size between species and recent work has also shown that it varies dramatically in gene content between species, such that there is no single gene shared between all GRCs sequenced (25 species total; Ruiz-Ruano et al., 2025; Torgasheva et al., 2019). This is particularly notable in birds, as unlike sciarids, the single GRC within each species is thought to originate from one chromosome. Additionally, GRCs do not seem to be particularly enriched in TEs, although there does seem to be some specific TE enrichment and also satellite expansion in some GRCs (Ruiz-Ruano et al., 2025). In fact, satellite evolution in GRCs seems like it might be fascinating. In both birds with GRCs and in sciarids, there is evidence that the satellite composition is largely distinct between GRCs and core chromosomes. This extends to the putative centromeric satellite in Sciarids (with the notable exception of *B. impatiens* which we believe is a recent turnover) and also in several nightingale species (Rídl et al., 2025). This difference between satellite composition in core chromosomes and GRCs in conjunction with the differences in chromosome behaviour over development (i.e. restriction to the germline and noncanonical transmission in meiosis), seems like it is unlikely to be coincidental. Therefore, GRCs are emerging as chromosomes that evolve in an extraordinarily dynamic fashion compared to the core chromosomes, and maintain a distinct satellite profile from the core genome.

We are now beginning to learn more about the nature of germline restricted chromosomes. So far the evidence suggests that these are in fact a distinct chromosome type: which likely evolves due to its germline restricted and initially redundant nature, as well as potentially selfishly, as transmission of these chromosomes occurs different than other chromosomes in the genome. Through the production of chromosome level genomes for the GRCs in sciarids, we have uncovered a number of important findings which will help with further studies of GRC evolution and nature in sciarids, specifically regarding the mechanism of GRC elimination, GRC gene evolution, and GRC architecture and evolution over time. This is an important stepping stone for future work on this clade.

## Methods

### Fly culture maintenance

The *B. coprophila* Holo2 stock was collected by Metz (Metz, 1925) and obtained through Susan Gerbi in 2017. The *B. impatiens* MIT line was obtained through Yukiko Yamashita in 2022, and the *L. ingenua* EDI line was obtained from Laura Ross in 2023. *Bradysia coprophila* and *B. impatiens* are monogenic (i.e. a female produces either male or female offspring, but not both sexes), while *L. ingenua* is digenic. We maintained flies in 30mL plastic vials, filled with 10 mL bacteriological agar. To reduce genetic diversity in the samples we sequenced, we mated siblings or near relatives, adding one female or one male into each vial, and raised the offspring produced. We fed larvae by sprinkling a mixture of powdered oat straw, mushrooms, spinach, and brewers yeast (Gerbi, 2024) into vials twice a week.

### Sample dissection, gDNA extraction, library preparation and sequencing

We tried to reduce genetic diversity among pooled individuals by preferentially sequencing only individuals from the same vial, or only individuals from vials in which we knew the parents were related. We carried out the testes dissections under a stereomicroscope on a clean microscope slide, dissecting each male in a drop of PBS. We dissected the testes by holding the male with a pin and gently pulling the claspers away from the body with another pin until the testes and male reproductive organs detached from the body, then we severed the vas deferens from the ejaculatory sac, transferring the testes and vas efferens into a 1.5 ml eppendorf tubes for sequencing.

For long read sequencing we pooled 13-21 testes frozen on dry ice per species. The testes were homogenised using a PowerMasher II tissue disruptor (Denton et al., 2023). DNA was extracted using a modified Qiagen MagAttract HMW DNA protocol and automated on a ThermoFisher KingFisher Apex (Oatley et al., 2023a). Due to the low DNA yield obtained from these samples, DNA was prepared for PacBio HiFi sequencing following the Ultra-low input library preparation. This method requires DNA fragments in the range of 8-10 kb. DNA was fragmented using a centrifuge-mediated method, following the Covaris g-tube adapted protocol (Oatley et al., 2023c). After fragmentation, the DNA was size selected using SPRI (solid-phase reversible immobilisation) beads and automated on the Kingfisher APEX (Oatley et al., 2023b). The resultant DNA was assessed by the Qubit fluorometric system, using the Qubit dsDNA High Sensitive Assay kit and Nanodrop spectrophotometric to evaluate concentration and purity, respectively. A Femto Pulse was used to assess fragment size distribution. Ultra-low-input (ULI) libraries were prepared using the PacBio SMRTbell® Express Template Prep Kit 2.0 and gDNA Sample Amplification Kit and sequenced on a Revio instrument (Pacific Biosciences) on a Revio 25M SMRT cell.

For Hi-C we used 30-41 pooled testes frozen on dry ice to generate libraries using Arima-HiC v2 kit (Arima Genomics). Following the manufacturer’s instructions, tissue was fixed and DNA crosslinked using TC buffer to a final formaldehyde concentration of 2%. The tissue was homogenised using the Diagnocine Power Masher-II. Crosslinked DNA was digested with a restriction enzyme master mix, biotinylated, and ligated. Clean-up was performed with SPRISelect beads before library preparation. Biotinylated DNA constructs were fragmented using a Covaris E220 sonicator and size selected to 400–600 bp using SPRISelect beads. DNA was enriched with Arima-HiC v2 kit Enrichment beads. End repair, A-tailing, and adapter ligation were carried out with the NEBNext Ultra II DNA Library Prep Kit (New England Biolabs), following a modified protocol where library preparation occurs while DNA remains bound to the Enrichment beads. Library amplification was performed using KAPA HiFi HotStart mix and a custom Unique Dual Index (UDI) barcode set (Integrated DNA Technologies). Depending on sample concentration and biotinylation percentage determined at the crosslinking stage, libraries were amplified with 10 to 16 PCR cycles. Post-PCR clean-up was performed with SPRISelect beads. Libraries were quantified using the AccuClear Ultra High Sensitivity dsDNA Standards Assay Kit (Biotium) and a FLUOstar Omega plate reader (BMG Labtech). Prior to sequencing, libraries were normalised to 10 ng/μL. Sequencing was performed using paired-end 150 bp reads on the Illumina NovaSeq 6000.

### Genome assembly

Prior to the assembly of the PacBio HiFi reads, we generated a database of *k*-mer counts (*k* = 31) with FastK (https://github.com/thegenemyers/FASTK) for each of the Sciaridae species, and visualised *k*-mer spectrum using GenomeScope package (Jenike et al., 2025; Ranallo-Benavidez et al., 2020; **Supplementary Figure 15**). We then assembled the PacBio HiFi reads into contigs with hifiasm (v.0.19.8-r603; Cheng et al. 2022) with the –primary option, and removed haplotypic duplications with purge_dups (v.1.2.5; Guan et al. 2020). We mapped the Hi-C read to the contigs with bwa-mem2 (v2.2.1; Vasimuddin et al., 2019), and joined them into scaffolds with YaHS (v1.2a.2; Zhou et al. 2023) using the --break option. We evaluated the completeness and contiguity of the genome assemblies using gfastats (Formenti et al., 2022), MERQURY.FK (Rhie et al., 2020) and BUSCO (v5.7.1; Waterhouse et al. 2018) in miniprot mode with the diptera_odb12 lineage. In addition to the nuclear genome, we identified and assembled the mitochondrial genome using MitoHiFi (v3; Uliano-Silva et al., 2023).

### Identification of the GRCs and genome curation

To assign scaffolds to chromosomes we applied the *k*-mer based technique described in Hodson et al. (2022). We used the kat comp command (Kmer Analysis Toolkit v2.4.2; Mapleson et al., 2017) to generate 2D *k*-mer spectra comparing 27-mer compositions between the germline PacBio library and publicly available non-germline libraries for the same species. We compared 1. Short read somatic libraries from PRJEB44837 for *B. coprophila* (Hodson et al., 2022), 2. Full body PacBio sequencing of a *B. impatiens* female from the same laboratory colony, 3. Full body short read sequencing data from PRJNA1109384 for *L. ingenua* (Baird et al., 2025; **Supplementary Figure 16**). We conducted the manual curation of the assemblies primarily in PretextView (https://github.com/sanger-tol/PretextView), taking scaffold assignments from above into account. Visual inspection and correction of the scaffolds was performed as described by (Howe et al., 2021). The curation process is documented at https://gitlab.com/wtsi-grit/rapid-curation. For generating high resolution Hi-C contact maps of the final assemblies, we used PretextSnapshot (https://github.com/sanger-tol/PretextSnapshot; see **Supplementary Figure 17** for Hi-C maps).

### Synteny analysis between chromosomes

To identify syntenic regions between chromosomes, we aligned pairs of genomes to each other using FastGA v1.1 with default parameters (https://github.com/thegenemyers/FASTGA). To retain only one-to-one relationships, we applied the ALNchain function from the FastGA toolkit. The resulting alignments were then converted into PAF format using the ALNtoPAF function and visualised with Oxford dotplots using custom R scripts.

### Gene prediction

Prior to performing gene annotation of the genomes, we identified transposable elements in each genome *de novo* with the EarlGrey TE annotation pipeline v3.0 using default parameters. The resulting species-specific repeat libraries were used as an input to RepeatMasker v4.1.5 for detection and soft-masking of repetitive sequences in the corresponding genomes. To predict protein-coding genes in Sciaridae genomes, we first split assemblies into the core and GRC sequences for each species and ran two rounds of Braker (version 2 or 3 depending on the evidence data) on them separately. For the first round, we used the Arthropoda protein database distributed with Braker, and annotated protein sequences of *Contarinia nasturtii* (GCF_009176525.2) as evidence for gene predictions for all genomes. Moreover, we added publicly available RNA-Seq data to the first round for *B. coprophila* (Kyriacou et al., 2025) and *L. ingenua* (Baird et al., 2025). Fly testes samples represent a mixture of somatic and germline tissue, so there is little evidence that genes in the GRCs are expressed in this tissue in adults. Therefore, to refine gene predictions in the GRCs, we ran a second round of Braker on these chromosomes, adding the core-derived protein sequences from the first round as extended support. We also used core- and GRC-derived protein sequences from *B. coprophila* and *L. ingenua* to enhance gene predictions in *B. impatiens*, due to the absence of RNA-Seq data for this species.

### BUSCO analysis of GRC gene origins and conservation between GRCs

We wanted to understand more about how the GRCs evolved and how similar they are to each other. To do this, we made use of conserved single copy orthologs (BUSCO) genes on the GRCs and core genomes of the three species we sequences, as well as in the publicly available genomes of several Sciaroidea species and an Anisopodidae outgroup species. A list of the species used for phylogenetic analyses is listed along with accession numbers is available in **Supplementary Table 2**. We ran BUSCO on each genome with parameter--metaeuk, and using database diptera_odb12 (Tegenfeldt et al., 2025). We analysed BUSCOs present in at least 60% of species, and for species other than the three that we sequenced (for which we expect a high number of duplicated BUSCOs given that the GRC genes almost always have a copy in the core genome) we retained the longest orf only for any duplicated BUSCOs. We then made amino acid alignments for each BUSCO gene with MAFFT linsi (v7.525; Katoh and Standley, 2013) using default settings. We generated gene trees for each BUSCO in IQtree (v2; Minh et al., 2020) the parameters -alrt 1000 -bb 1000. We analysed only gene trees for which the Cecidomyiidae and Sciaridae sequences (excepting the GRC sequences) were both monophyletic. Therefore, we ran a custom R script which summarized whether cecidomyiid and sciarid clades are monophyletic for each BUSCO tree, and when this was the case, which clade GRC genes fell into and what species was the nearest neighbour. We found that 19 BUSCOs on the GRCs in *L. ingenua* were located on unplaced scaffolds, and since we wanted to understand the evolutionary dynamics for each GRC, we removed these genes from the analysis. We also assessed whether the same BUSCO gene was found on different GRCs with the same origin, to get a sense of the conservation of gene architecture on the GRCs, and how BUSCOs were spatially organized on each GRC. We then ran ASTRAL-Pro-3 (Zhang et al., 2025, 2020) grouping core genes based on species and GRCs based on GRC chromosome and origin (i.e. for each GRC we grouped the genes belonging to the Cecidomyiidae clade and Sciaridae clade separately). Although we acknowledge that this approach may give less fine scale resolution for the phylogenetic placement of blocks of genes moving onto the GRCs from the core genome, we wanted to maximize the data we retained within Sciaridae, by not removing genetrees with unusual placements of GRC genes or with low bootstrap support.

### Tandem repeat annotation and prediction of centromeric regions

We identified tandem repeats in the three Sciaridae genomes using the Tandem Repeat Annotation and Structural Hierarchy (TRASH) pipeline (Wlodzimierz et al., 2023) with default parameters. We further calculated the density of these repeats and GC content in 50-kbp windows across each chromosome, marking all the regions where a drop in GC content overlapped with a peak in tandem repeats as putative centromeres. We further assessed these regions against the Hi-C contact maps to see which of them overlap with the off-diagonal signals of spatial centromere clustering observed for some of the chromosomes. When centromere positions were less clear or the off-diagonal Hi-C signal was not present, we examined satellite content across all candidate regions and prioritized regions that contained satellite families shared among the same chromosome types as primary centromere candidates (e.g., between GRCs or among core chromosomes within a genome).

To compare centromere-specific satellite DNA sequences across chromosomes within the same genome, we extracted copies of the most abundant satellite repeats from the annotated centromeres and calculated *k*-mer–based Jaccard similarity scores (*k* = 6) for all pairwise combinations using a custom R script. We clustered the similarity scores with the hierarchical cluster analysis and visualized the results using the pheatmap R package (v1.0.13). To confirm whether GRC- or core-specific satellite DNAs were absent in other chromosome groups, we extracted all copies of the satellites from the centromeric regions from the TRASH raw file. We then generated a consensus sequence from a multiple sequence alignment produced with MAFFT (v7.525) and searched it against the rest of the genome with BLASTn (v2.15.0) using exact length match and more than 96% identity as cutoffs.

### Feature distribution along chromosomes

To investigate the distribution of various genomic features, such as transposable elements and protein-coding genes, relative to the centromere position on each chromosome, we first divided each chromosome into arms based on the centromere coordinates predicted above. Chromosome arms representing less than 35% of the total chromosome length were excluded from further analysis. We then used the bed_makewindows function from the valr R package (v0.8.3; Riemondy et al., 2017) to split each chromosome arm into 100 equally sized windows, and intersected these windows with the genomic features to calculate the per-window coverage of each feature independently using the bed_coverage function.

### Identification of the controlling element

To search for the controlling element in *B. coprophila*, we aligned nucleotide sequences of GRCs to the X chromosome with FastGA as described above and extracted one-to-one alignments that were only shared by the short arm of the X chromosome (SUPER_X:65600000-69053365) and the GRCs, ending up with two synteny blocks found in all three. To confirm that these alignments are unique to these regions, we extracted their nucleotide sequences using the getfasta command from bedtools (v2.31.1) and compared them against the rest of the *B. coprophila* genome using BLASTn (v2.15.0). We also looked at whether any tandem repeats predicted by TRASH are present only in the short arm of the X chromosome and GRCs. For that, we extracted all repeat sequences from the raw file generated by TRASH for the short arm of the chromosome X, clustered them by length and predicted consensus sequences using the method described above. These sequences were again used to search against the rest of the genome with BLASTn (v2.15.0). To anchor these sequences to regions of the X chromosome known to be associated with the controlling element, we blasted sequences identified in the controlling element from Escribá et al. (2011), and Burke et al. (1993). This allowed us to identify the H1 region proximal to the X chromosome centromere and the H2/H3 region, which is expected to contain the controlling element in *B. coprophila* (Crouse 1977; Crouse et al., 1977).

### Inferring orthology and purifying selection

We downloaded genomes and annotations for *D. melanogaster* (GCF_000001215.4), *A. aphidimyza*, *O. robiniae*, *C. nasturtii*, and *P. hygida* (accessions in **Supplementary Table 2;** (Hoskins et al., 2015; Huang et al., 2024; Mori et al., 2021; Shen et al., 2024; Trinca et al., 2023) to include with the aforementioned sciarid core and germline-restricted chromosome de-novo annotations in orthology and selection inference. We used AGAT (v1.4.1) to select the longest isoforms of complete proteins and extracted the nucleotide coding sequences and translated proteins from each genome. We then inferred orthology using Orthofinder (v3.0.1.b1) including the multiple sequence alignment (-M msa) option to infer gene trees. We stratified orthogroups by GRC gene origin using a custom script (https://github.com/Obscuromics/Fly-Germ-Line-Chromosomes/blob/main/scripts/nwk2nwk_renamed_tips_by_origin.py) and proceeded using all successful classifications (5,759/24,314 orthogroups). Orthology inference produced very few single-copy ortholog groups between the chosen proteomes (82). None of the GRCs’ proteomes constitute an approximation of the core. When specifically considering orthogroups with single representatives from all present core genomes, each GRC appears to lack a representative in a large number of cases, which is not true of the core genomes each considered in turn. Hence in absence of clear encompassing orthogroup selection criteria, we classified 5674 orthogroups containing at least one orthologous core and GRC gene pair from either Sciaridae or Cecidomyiidae. We performed pre-alignment quality control on the aggregated coding sequences of each orthogroup using Prequal (v1.02) followed by codon-aware alignment with macsev2 (v2.07). Using these alignments and OrthoFinder’s gene trees we pruned filtered-out genes using Biopython, and inferred purifying selection using codeml (v4.10.7). We ran codeml with a free-parameter model to infer dn/ds rates per branch in each orthogroup using the following settings: ‘noisy=9, verbose=1, runmode=0, seqtype=1, CodonFreq=2, clock=0, model=1, NSsites=0, icode=0, fix_kappa=0, kappa,=2, fix_omega=0, omega=0.5, and cleandata=1’.

For all of the analysed orthogroups, we identified whether each branch led to only GRC genes, Sciaridae core genes, Cecidomyiidae genes, or other species. We compared the dn/ds of branches for each of these categories (**Supplementary Figure 18)**. Then we did a more specific comparison where we took only genes with one copy on the GRC within a species and one copy on the core genome of the same species (4133 genes total). We classified whether the GRC homolog fell within the Cecidomyiidae (1992 genes) or Sciaridae clade (1293 genes) in the phylogeny (or was unplacable). For genes in the Cecidomyiidae clade, we compared the dN/dS of the GRC branch (or branches if several GRC genes formed a clade), to the Sciaridae clade branches from the core gene of the same species (as the focal GRC gene) to the crown of the Sciaridae lineage. The expectation was that since GRCs evolved at the base of the Sciaridae clade, this would represent a reasonable comparison. For GRC genes in the Sciaridae clade, we compared the dN/dS of the GRC branch to the core branch until they met within the Sciaidae (See **Supplementary Figure 7** for explanatory schematic). The rationale behind this analysis was to see whether the GRC genes were evolving in a different way as their core homolog over a similar time period. We conducted a paired t-test for Cecidomyiidae and Sciaridae origin genes separately using the Welsh approximation to estimate variance. We also compared whether the GRC branch dN/dS differed from the gene tree as a whole, by running codeml with two omegas: one for the GRC-gene terminal branch (foreground) and one for the entire tree (background). This analysis was done only when the terminal branch was at least 0.1 long, as with lower branch lengths dN/dS estimates become too noisy (**Supplementary Figure 8**).

### Analysis of recently duplicated syntenic blocks onto the GRCs

We also looked at how GRC genes are evolving over time by comparing the duplicated syntenic blocks of Sciarid origin on the GRCs to their core counterparts. We considered syntenic blocks on any core chromosome and GRC which were over 200 kbp (on the core chromosome) with at least 10 paralog pairs (with one gene copy in the core genome and one gene copy in the syntenic block on the GRC and no other homologs in the genome). This left us with 37 blocks for *B. impatiens,* all between either autosome 1 or the X chromosome and the GRC and up to ∼1.5 Mbp, and 4 blocks for both *B. coprophila* and *L. ingenua,* between autosome 2 or the X chromosome and GRC2 (for *B. coprophila*) and autosome 3 and GRC1 or autosome 1 and GRC2 (for *L. ingenua)* and up to ∼400 kbp for both species (**Supplementary Table 3**). We assessed how genes on the GRC were evolving over time compared to their core counterparts, by comparing the number of genes on each syntenic block, whether genes on the GRC portion of the syntenic block were duplicated or missing compared to the core chromosome portion, and for missing genes, whether there was any remnant of the gene remaining in the syntenic block by conducting blastn searches of the remaining gene against the genome (**Supplementary Table 4**). We compared the length and exon number of the paralog pairs and also assessed the dS of as many genes on each syntenic block as possible, to get an idea of the divergence time since the duplication onto the GRC happened. We did this for all gene pairs in which one gene copy was on a GRC and one in the core genome and for which we could determine the gene origin (**Supplementary Figure 19**), however, we focused our main analysis on genes in syntenic blocks since we have a more clear idea of the genomic architecture (i.e. gene order along blocks) of those genes so we could assess whether GRC genes had been lost over time.

## Data, Materials, and Software Availability

Sequence data and assemblies used in this study are publically available: *Bradysia coprophila* (GCA_965233685.1; SRA: ERS15411730), *Bradysia impatiens* (GCA_965233325.1; SRA: ERS15411732), *Lycoriella ingenua* (GCA_965233655.1; SRA: ERS15411740). Genomes for *Diadocidia ferruginosa, Acnemia nitidicollis, Coelosia flava, Bradysia nitidicollis, Bradysia ocellaris, Corynoptera forcipata, Phytosciara flavipes*, and *Schwenckfeldina carbonaria*, used in phylogenetic analyses will be available upon publication. Scripts associated with the analyses in this study can be found at https://github.com/Obscuromics/Fly-Germ-Line-Chromosomes.

## Acknowledgements

We would like to thank Laura Ross and Rob Baird for helpful comments throughout this research and for providing the *L. ingenua* colony. We would also like to thank Yukiko Yamashita and Susan Gerbi for providing the *B. impatiens* and *B. coprophila* colonies. We would like to thank the Genome Engine of the Tree of Life programme for help with generating the reference genomes. CNH is funded by a Royal Society Newton International Fellowship (NIF\R1\232397). This research was funded in whole, or in part, by the Wellcome Trust 220540/Z/20/A. For the purpose of Open Access, the author has applied a CC BY public copyright licence to any Author Accepted Manuscript version arising from this submission.

## Notes

### Competing Interest Statement

The authors have declared no competing interest.

### Summary of Updates

Added figure 3 and revised text

## References

Baduel, P., Bray, S., Vallejo-Marin, M., Kolář, F., Yant, L., 2018. The “Polyploid Hop”: Shifting Challenges and Opportunities Over the Evolutionary Lifespan of Genome Duplications. Front. Ecol. Evol. 6. 10.3389/fevo.2018.00117

Bauer, H., Beermann, W., 1952. Der Chromosomencyclus der Orthocladiinen (Nematocera, Diptera). Z. Für Naturforschung B 7, 557–563. 10.1515/znb-1952-9-1013

Beadle, G.W., 1932. The Relation of Crossing over to Chromosome Association in Zea-Euchlaena Hybrids. Genetics 17, 481–501. 10.1093/genetics/17.4.481

Bénitière, F., Arnaiz, O., Penel, S., Duharcourt, S., Meyer, E., Sperling, L., Duret, L., 2026. Breaking the species barrier: recurrent genomic introgressions from very distant lineages in a ciliate. 10.64898/2026.02.20.707023

Burke, W.D., Eickbush, D.G., Xiong, Y., Jakubczak, J., Eickbush, T.H., 1993. Sequence relationship of retrotransposable elements R1 and R2 within and between divergent insect species. Mol. Biol. Evol. 10, 163–185. 10.1093/oxfordjournals.molbev.a039990

Carson, H.L., 1944. An analysis of natural chromosome variability in Sciara impatiens Johannsen. J. Morphol. 75, 11–59. 10.1002/jmor.1050750103

Charlesworth, B., Harvey, P.H., Charlesworth, Brian, Charlesworth, D., 2000. The degeneration of Y chromosomes. Philos. Trans. R. Soc. Lond. B. Biol. Sci. 355, 1563–1572. 10.1098/rstb.2000.0717

Cheng, H., Jarvis, E.D., Fedrigo, O., Koepfli, K.-P., Urban, L., Gemmell, N.J., Li, H., 2022. Haplotype-resolved assembly of diploid genomes without parental data. Nat. Biotechnol. 40, 1332–1335. 10.1038/s41587-022-01261-x

Crane, Y.M., Crane, C.F., Cambron, S.E., Springmeyer, L.J., Schemerhorn, B.J., 2023. Molecular characterization of eliminated chromosomes in Hessian fly (Mayetiola destructor (Say)). Chromosome Res. Int. J. Mol. Supramol. Evol. Asp. Chromosome Biol. 31, 3. 10.1007/s10577-023-09718-8

Crouse, H.V., 1979. X heterochromatin subdivision and cytogenetic analysis in Sciara coprophila (diptera, sciaridae). Chromosoma 74, 219–239. 10.1007/BF00292274

Crouse, H.V., 1977. X heterochromatin subdivision and cytogenetic analysis in Sciara coprophila (Diptera, Sciaridae). Chromosoma 63, 39–55. 10.1007/BF00292941

Crouse, H.V., 1943. Translocations in Sciara : their bearing on chromosome behavior and sex determination.

Crouse, H.V., Brown, A., Mumford, B.C., 1971. L-chromosome inheritance and the problem of chromosome “imprinting” in Sciara (Sciaridae, Diptera). Chromosoma 34, 324–339. 10.1007/BF00286156

Crouse, H.V., Gerbi, S.A., Liang, C.M., Magnus, L., Mercer, I.M., 1977. Localization of ribosomal DNA within the proximal X heterochromatin of Sciara coprophila (Diptera, Sciaridae). Chromosoma 64, 305–318. 10.1007/BF00294938

de Saint Phalle, B., Sullivan, W., 1996. Incomplete sister chromatid separation is the mechanism of programmed chromosome elimination during early Sciara coprophila embryogenesis. Dev. Camb. Engl. 122, 3775–3784. 10.1242/dev.122.12.3775

Denton, A., Oatley, G., Cornwell, C., Quail, M., Howard, C., 2023. Sanger Tree of Life Sample Homogenisation: PowerMash.

Du Bois, A.M., 1933. Chromosome behavior during cleavage in the eggs of Sciara coprophila (Diptera) in the relation to the problem of sex determination. Z. Für Zellforsch. Mikrosk. Anat. 19, 595–614. 10.1007/BF00393361

DuBois, A.M., 1932. Elimination of Chromosomes during Cleavage in the Eggs of Sciara (Diptera)1. Proc. Natl. Acad. Sci. 18, 352–356. 10.1073/pnas.18.5.352

Escribá, M.C., Greciano, P.G., Méndez-Lago, M., de Pablos, B., Trifonov, V.A., Ferguson-Smith, M.A., Goday, C., Villasante, A., 2011. Molecular and cytological characterization of repetitive DNA sequences from the centromeric heterochromatin of Sciara coprophila. Chromosoma 120, 387–397. 10.1007/s00412-011-0320-2

Formenti, G., Abueg, L., Brajuka, A., Brajuka, N., Gallardo-Alba, C., Giani, A., Fedrigo, O., Jarvis, E.D., 2022. Gfastats: conversion, evaluation and manipulation of genome sequences using assembly graphs. Bioinformatics 38, 4214–4216. 10.1093/bioinformatics/btac460

Gerbi, S.A., 2024. Laboratory Maintenance of the Lower Dipteran Fly Bradysia (Sciara) coprophila: A New/Old Emerging Model Organism. J. Vis. Exp. JoVE e66751. 10.3791/66751

Gerbi, S.A., 2022. Non-random chromosome segregation and chromosome eliminations in the fly Bradysia (Sciara). Chromosome Res. Int. J. Mol. Supramol. Evol. Asp. Chromosome Biol. 30, 273–288. 10.1007/s10577-022-09701-9

Geyer-Duszyńska, I., 1959. Experimental research on chromosome elimination in cecidomyidae (Diptera). J. Exp. Zool. 141, 391–447. 10.1002/jez.1401410302

Gruzova, M., Batalova, F., 1993. Oogenesis and meiotic divisions of predatory gall midge, *Aphidoletes aphidimyza Rond.* (Diptera : Cecidomyiidae). Int. J. Insect Morphol. Embryol., Cuticle, Stridulatory and Hearing Organs, Ovarioles and Oogenesis, Egg Chorion, Spermatozoa, and Midgut Cell Junctions 22, 315–334. 10.1016/0020-7322(93)90017-U

Guan, D., McCarthy, S.A., Wood, J., Howe, K., Wang, Y., Durbin, R., 2020. Identifying and removing haplotypic duplication in primary genome assemblies. Bioinformatics 36, 2896–2898. 10.1093/bioinformatics/btaa025

Haig, D., 1993. The evolution of unusual chromosomal systems in sciarid flies: intragenomic conflict and the sex ratio. J. Evol. Biol. 6, 249–261. 10.1046/j.1420-9101.1993.6020249.x

Hodson, C.N., Jaron, K.S., Gerbi, S., Ross, L., 2022. Gene-rich germline-restricted chromosomes in black-winged fungus gnats evolved through hybridization. PLOS Biol. 20, e3001559. 10.1371/journal.pbio.3001559

Hodson, C.N., Ross, L., 2021. Evolutionary Perspectives on Germline-Restricted Chromosomes in Flies (Diptera). Genome Biol. Evol. 13, evab072. 10.1093/gbe/evab072

Hoskins, R.A., Carlson, J.W., Wan, K.H., Park, S., Mendez, I., Galle, S.E., Booth, B.W., Pfeiffer, B.D., George, R.A., Svirskas, R., Krzywinski, M., Schein, J., Accardo, M.C., Damia, E., Messina, G., Méndez-Lago, M., de Pablos, B., Demakova, O.V., Andreyeva, E.N., Boldyreva, L.V., Marra, M., Carvalho, A.B., Dimitri, P., Villasante, A., Zhimulev, I.F., Rubin, G.M., Karpen, G.H., Celniker, S.E., 2015. The Release 6 reference sequence of the Drosophila melanogaster genome. Genome Res. 25, 445–458. 10.1101/gr.185579.114

Howe, K., Chow, W., Collins, J., Pelan, S., Pointon, D.-L., Sims, Y., Torrance, J., Tracey, A., Wood, J., 2021. Significantly improving the quality of genome assemblies through curation. GigaScience 10, giaa153. 10.1093/gigascience/giaa153

Huang, L., Wang, L., Sun, H.-Q., Huai, W.-X., Lin, R.-Z., Wei, S.-J., Yao, Y.-X., 2024. The chromosome-level genome assembly and annotation of an invasive forest pest Obolodiplosis robiniae. Sci. Data 11, 1227. 10.1038/s41597-024-04037-x

Jenike, K.M., Campos-Domínguez, L., Boddé, M., Cerca, J., Hodson, C.N., Schatz, M.C., Jaron, K.S., 2025. k-mer approaches for biodiversity genomics. Genome Res. 35, 219–230. 10.1101/gr.279452.124

Johnson Pokorná, M., Reifová, R., 2021. Evolution of B Chromosomes: From Dispensable Parasitic Chromosomes to Essential Genomic Players. Front. Genet. 12. 10.3389/fgene.2021.727570

Kahle, W., 1908. Die Paedogenesis der Decidomyiden, Zoologica ;Hft. 55. E. Schweizerbart, Stuttgart.

Katoh, K., Standley, D.M., 2013. MAFFT Multiple Sequence Alignment Software Version 7: Improvements in Performance and Usability. Mol. Biol. Evol. 30, 772–780. 10.1093/molbev/mst010

Kerrebrock, A., Flynn, J.M., Baird, R.B., Hodson, C.N., Ross, L., Yamashita, Y.M., 2025. Distinct satellite DNA composition between core and germline restricted chromosomes in Bradysia (Sciara) coprophila. G3 Bethesda Md 15, jkaf155. 10.1093/g3journal/jkaf155

Kinsella, C.M., Ruiz-Ruano, F.J., Dion-Côté, A.-M., Charles, A.J., Gossmann, T.I., Cabrero, J., Kappei, D., Hemmings, N., Simons, M.J.P., Camacho, J.P.M., Forstmeier, W., Suh, A., 2019. Programmed DNA elimination of germline development genes in songbirds. Nat. Commun. 10, 5468. 10.1038/s41467-019-13427-4

Kyriacou, R.G., Herbette, M., Baird, R.B., Monteith, K.M., Ross, L., Hodson, C.N., 2025. Evidence for Transcription and Horizontal Gene Transfer in Dipteran Germline-Restricted Chromosomes. 10.64898/2025.12.11.693641

Mapleson, D., Garcia Accinelli, G., Kettleborough, G., Wright, J., Clavijo, B.J., 2017. KAT: a K-mer analysis toolkit to quality control NGS datasets and genome assemblies. Bioinforma. Oxf. Engl. 33, 574–576. 10.1093/bioinformatics/btw663

Meier, R., Lawniczak, M.K.N., Srivathsan, A., 2025. Illuminating Entomological Dark Matter with DNA Barcodes in an Era of Insect Decline, Deep Learning, and Genomics. Annu. Rev. Entomol. 70, 185–204. 10.1146/annurev-ento-040124-014001

Metz, Charles W., 1938. Chromosome Behavior, Inheritance and Sex Determination in Sciara. Am. Nat. 72, 485–520. 10.1086/280803

Metz, C. W., 1938. Chromosome Behavior, Inheritance and Sex Determination in Sciara. Am. Nat. 72, 485–520.

Metz, C.W., 1926. Genetic Evidence of a Selective Segregation of Chromosomes in Sciara (Diptera). Proc. Natl. Acad. Sci. 12, 690–692. 10.1073/pnas.12.12.690

Metz, C.W., 1925. Chromosomes and Sex in Sciara. Science 61, 212–214. 10.1126/science.61.1573.212

Metz, C.W., Moses, M.S., Hoppe, E.N., 1926. Chromosome behavior in Sciara (Diptera). I: Chromosome behavior in the spermatocyte divisions. Z. Für Indukt. Abstammmungs Vererbungslehre 42, 237–270. 10.1007/BF01741805

Minh, B.Q., Schmidt, H.A., Chernomor, O., Schrempf, D., Woodhams, M.D., von Haeseler, A., Lanfear, R., 2020. IQ-TREE 2: New Models and Efficient Methods for Phylogenetic Inference in the Genomic Era. Mol. Biol. Evol. 37, 1530–1534. 10.1093/molbev/msaa015

Mori, B.A., Coutu, C., Chen, Y.H., Campbell, E.O., Dupuis, J.R., Erlandson, M.A., Hegedus, D.D., 2021. De Novo Whole-Genome Assembly of the Swede Midge (Contarinia nasturtii), a Specialist of Brassicaceae, Using Linked-Read Sequencing. Genome Biol. Evol. 13, evab036. 10.1093/gbe/evab036

Naish, M., Alonge, M., Wlodzimierz, P., Tock, A.J., Abramson, B.W., Schmücker, A., Mandáková, T., Jamge, B., Lambing, C., Kuo, P., Yelina, N., Hartwick, N., Colt, K., Smith, L.M., Ton, J., Kakutani, T., Martienssen, R.A., Schneeberger, K., Lysak, M.A., Berger, F., Bousios, A., Michael, T.P., Schatz, M.C., Henderson, I.R., 2021. The genetic and epigenetic landscape of the Arabidopsis centromeres. Science 374, eabi7489. 10.1126/science.abi7489

Nakai, Y., Kubota, S., Kohno, S., 1991. Chromatin diminution and chromosome elimination in four Japanese hagfish species. Cytogenet. Cell Genet. 56, 196–198. 10.1159/000133087

Oatley, G., Denton, A., Howard, C., 2023a. Sanger Tree of Life HMW DNA Extraction: Automated MagAttract v.2.

Oatley, G., Sampaio, F., Howard, C., 2023b. Sanger Tree of Life Fragmented DNA clean up: Automated SPRI.

Oatley, G., Sampaio, F., Kitchin, L., Amaral, R.J.V. do, Howard, C., 2023c. Sanger Tree of Life HMW DNA Fragmentation: Covaris g-TUBE for ULI PacBio.

Pei, Y., Forstmeier, W., Ruiz-Ruano, F.J., Mueller, J.C., Cabrero, J., Camacho, J.P.M., Alché, J.D., Franke, A., Hoeppner, M., Börno, S., Gessara, I., Hertel, M., Teltscher, K., Knief, U., Suh, A., Kempenaers, B., 2022. Occasional paternal inheritance of the germline-restricted chromosome in songbirds. Proc. Natl. Acad. Sci. 119, e2103960119. 10.1073/pnas.2103960119

Pigozzi, M.I., Solari, A.J., 2005. The germ-line-restricted chromosome in the zebra finch: recombination in females and elimination in males. Chromosoma 114, 403–409. 10.1007/s00412-005-0025-5

Pigozzi, M.I., Solari, A.J., 1998. Germ cell restriction and regular transmission of an accessory chromosome that mimics a sex body in the zebra finch, Taeniopygia guttata. Chromosome Res. Int. J. Mol. Supramol. Evol. Asp. Chromosome Biol. 6, 105–113. 10.1023/a:1009234912307

Ranallo-Benavidez, T.R., Jaron, K.S., Schatz, M.C., 2020. GenomeScope 2.0 and Smudgeplot for reference-free profiling of polyploid genomes. Nat. Commun. 11, 1432. 10.1038/s41467-020-14998-3

Rhie, A., Walenz, B.P., Koren, S., Phillippy, A.M., 2020. Merqury: reference-free quality, completeness, and phasing assessment for genome assemblies. Genome Biol. 21, 245. 10.1186/s13059-020-02134-9

Rídl, J., Dedukh, D., Halenková, Z., Schlebusch, S.A., Beneš, V., Lopez, M.O., Osiejuk, T.S., Ruiz-Ruano, F.J., Suh, A., Albrecht, T., Reif, J., Reifová, R., 2025. Germline-restricted chromosome of songbirds has different centromere compared to regular chromosomes. Heredity 1–8. 10.1038/s41437-025-00779-5

Rieffel, S.M., Crouse, H.V., 1966. The elimination and differentiation of chromosomes in the germ line of sciara. Chromosoma 19, 231–276. 10.1007/BF00326917

Riemondy, K.A., Sheridan, R.M., Gillen, A., Yu, Y., Bennett, C.G., Hesselberth, J.R., 2017. valr: Reproducible genome interval analysis in R. F1000Research 6, 1025. 10.12688/f1000research.11997.1

Ruiz-Ruano, F.J., Schlebusch, S.A., Vontzou, N., Moreno, H., Biegler, M.T., Kutschera, V.E., Ekman, D., Borges, I., Pei, Y., Rossini, R., Albrecht, T., Boman, J., Borodin, P., Burri, R., Cain, K.E., Forstmeier, W., Frankl-Vilches, C., Gahr, M., Griffith, S.C., Hill, A.M., Irestedt, M., Joseph, L., Jønsson, K.A., Kawakami, T., Kempenaers, B., Malinovskaya, L., Mueller, J.C., Oliveira, E.H.C. de, Palacios-Gimenez, O.M., Palinauskas, V., Qvarnström, A., Reifova, R., Ridl, J., Segami, J.C., Tan, D.J.X., Torgasheva, A., Whibley, A., Suh, A., 2025. Programmed DNA elimination drives rapid genomic innovation in two thirds of all bird species. 10.1101/2025.07.16.664580

Schlebusch, S.A., Rídl, J., Poignet, M., Ruiz-Ruano, F.J., Reif, J., Pajer, P., Pačes, J., Albrecht, T., Suh, A., Reifová, R., 2023. Rapid gene content turnover on the germline-restricted chromosome in songbirds. Nat. Commun. 14, 4579. 10.1038/s41467-023-40308-8

Shen, X., Jin, J., Zhang, G., Yan, B., Yu, X., Wu, H., Yang, M., Zhang, F., 2024. The chromosome-level genome assembly of Aphidoletes aphidimyza Rondani (Diptera: Cecidomyiidae). Sci. Data 11, 785. 10.1038/s41597-024-03614-4

Smith, J.J., Timoshevskaya, N., Ye, C., Holt, C., Keinath, M.C., Parker, H.J., Cook, M.E., Hess, J.E., Narum, S.R., Lamanna, F., Kaessmann, H., Timoshevskiy, V.A., Waterbury, C.K.M., Saraceno, C., Wiedemann, L.M., Robb, S.M.C., Baker, C., Eichler, E.E., Hockman, D., Sauka-Spengler, T., Yandell, M., Krumlauf, R., Elgar, G., Amemiya, C.T., 2018. The sea lamprey germline genome provides insights into programmed genome rearrangement and vertebrate evolution. Nat. Genet. 50, 270–277. 10.1038/s41588-017-0036-1

Staiber, W., 2006. Chromosome elimination in germ line – soma differentiation of Acricotopus lucidus (Diptera, Chironomidae). Genome 49, 269–274. 10.1139/g05-103

Tegenfeldt, F., Kuznetsov, D., Manni, M., Berkeley, M., Zdobnov, E.M., Kriventseva, E.V., 2025. OrthoDB and BUSCO update: annotation of orthologs with wider sampling of genomes. Nucleic Acids Res. 53, D516–D522. 10.1093/nar/gkae987

Timoshevskiy, V.A., Herdy, J.R., Keinath, M.C., Smith, J.J., 2016. Cellular and Molecular Features of Developmentally Programmed Genome Rearrangement in a Vertebrate (Sea Lamprey: Petromyzon marinus). PLOS Genet. 12, e1006103. 10.1371/journal.pgen.1006103

Timoshevskiy, V.A., Timoshevskaya, N., Eşkut, K.I., Rajandran, K., Smith, J.J., 2025. Biparental inheritance of germline-specific chromosomes in the sea lamprey and their roles in oocytes. Proc. Natl. Acad. Sci. 122, e2421883122. 10.1073/pnas.2421883122

Torgasheva, A.A., Malinovskaya, L.P., Zadesenets, K.S., Karamysheva, T.V., Kizilova, E.A., Akberdina, E.A., Pristyazhnyuk, I.E., Shnaider, E.P., Volodkina, V.A., Saifitdinova, A.F., Galkina, S.A., Larkin, D.M., Rubtsov, N.B., Borodin, P.M., 2019. Germline-restricted chromosome (GRC) is widespread among songbirds. Proc. Natl. Acad. Sci. 116, 11845–11850. 10.1073/pnas.1817373116

Trinca, V., Carli, S., Uliana, J.V.C., Garbelotti, C.V., Mendes da Silva, M., Kunes, V., Meleiro, L.P., Brancini, G.T.P., Menzel, F., Andrioli, L.P.M., Torres, T.T., Ward, R.J., Monesi, N., 2023. Biocatalytic potential of Pseudolycoriella CAZymes (Sciaroidea, Diptera) in degrading plant and fungal cell wall polysaccharides. iScience 26, 106449. 10.1016/j.isci.2023.106449

Uliano-Silva, M., Ferreira, J.G.R.N., Krasheninnikova, K., Blaxter, M., Mieszkowska, N., Hall, N., Holland, P., Durbin, R., Richards, T., Kersey, P., Hollingsworth, P., Wilson, W., Twyford, A., Gaya, E., Lawniczak, M., Lewis, O., Broad, G., Martin, F., Hart, M., Barnes, I., Formenti, G., Abueg, L., Torrance, J., Myers, E.W., Durbin, R., Blaxter, M., McCarthy, S.A., Darwin Tree of Life Consortium, 2023. MitoHiFi: a python pipeline for mitochondrial genome assembly from PacBio high fidelity reads. BMC Bioinformatics 24, 288. 10.1186/s12859-023-05385-y

Urban, J.M., Gerbi, S.A., Spradling, A.C., 2025. Chromosome-scale scaffolds of the fungus gnat genome reveal multi-Mb-scale chromosome-folding interactions, centromeric enrichments of retrotransposons, and candidate telomere sequences. BMC Genomics 26, 443. 10.1186/s12864-025-11573-2

Vasimuddin, Md., Misra, S., Li, H., Aluru, S., 2019. Efficient Architecture-Aware Acceleration of BWA-MEM for Multicore Systems, in: 2019 IEEE International Parallel and Distributed Processing Symposium (IPDPS). Presented at the 2019 IEEE International Parallel and Distributed Processing Symposium (IPDPS), pp. 314–324. 10.1109/IPDPS.2019.00041

Vontzou, N., Pei, Y., Campo-Bes, I., Forstmeier, W., Hertel, M., Irimia, M., Kempenaers, B., Kuhn, S., Martin, K., Mueller, J.C., Teltscher, K., Mollbrink, A., Abalo, X., Biegler, M.T., Immler, S., Ruiz-Ruano, F.J., Suh, A., 2025. The germline-restricted chromosome orchestrates germ cell development in passerine birds. 10.64898/2025.12.02.691941

Vontzou, N., Pei, Y., Mueller, J.C., Reifová, R., Ruiz-Ruano, F.J., Schlebusch, S.A., Suh, A., 2023. Songbird germline-restricted chromosome as a potential arena of genetic conflicts. Curr. Opin. Genet. Dev. 83, 102113. 10.1016/j.gde.2023.102113

Wang, J., Davis, R.E., 2014. Programmed DNA elimination in multicellular organisms. Curr. Opin. Genet. Dev., Developmental mechanisms, patterning and evolution 27, 26–34. 10.1016/j.gde.2014.03.012

Waterhouse, R.M., Seppey, M., Simão, F.A., Manni, M., Ioannidis, P., Klioutchnikov, G., Kriventseva, E.V., Zdobnov, E.M., 2018. BUSCO Applications from Quality Assessments to Gene Prediction and Phylogenomics. Mol. Biol. Evol. 35, 543–548. 10.1093/molbev/msx319

White, M.J.D., 1973. Animal Cytology and Evolution. Cambridge University Press, Cambridge, Eng.

Wlodzimierz, P., Hong, M., Henderson, I.R., 2023. TRASH: Tandem Repeat Annotation and Structural Hierarchy. Bioinformatics 39, btad308. 10.1093/bioinformatics/btad308

Yasmin, T., Grayson, P., Docker, M.F., Good, S.V., 2022. Pervasive male-biased expression throughout the germline-specific regions of the sea lamprey genome supports key roles in sex differentiation and spermatogenesis. Commun. Biol. 5, 434. 10.1038/s42003-022-03375-z

Zhang, C., Nielsen, R., Mirarab, S., 2025. ASTER: A Package for Large-Scale Phylogenomic Reconstructions. Mol. Biol. Evol. 42, msaf172. 10.1093/molbev/msaf172

Zhang, C., Scornavacca, C., Molloy, E.K., Mirarab, S., 2020. ASTRAL-Pro: Quartet-Based Species-Tree Inference despite Paralogy. Mol. Biol. Evol. 37, 3292–3307. 10.1093/molbev/msaa139

Zhou, C., McCarthy, S.A., Durbin, R., 2023. YaHS: yet another Hi-C scaffolding tool. Bioinformatics 39, btac808. 10.1093/bioinformatics/btac808

