## Supplementary information for "Germline restricted chromosomes in dark winged fungus gnats show dynamic chromosome organisation despite strong purifying selection on genes"

**Supplementary Table 1.** Correspondence between the core chromosomes in our assembly and those from the recently published reference genome for *B. coprophila* (Urban et al. 2025), corresponding to the chromosome identification in this species by Crouse (1943). Note we followed the naming scheme used by the tree of life programme at Welcome Sanger Institute, with the largest chromosome labelled SUPER_1 etc.

| From literature (Urban et al., 2022; Couse 1943) | Our genome |
|---|---|
| Chromosome II | SUPER_3 |
| Chromosome III | SUPER_2 |
| Chromosome IV | SUPER_1 |
| Chromosome X | SUPER_X |

**Supplementary Table 2.** Species used for BUSCO analyses. Available reference assemblies within the Bibionomorpha clade are shown, several were excluded from analyses (indicated with asterisks) due to low proportion of complete BUSCOs (<60%). The assemblies we generated are indicated in bold.

| Species | Family | Accession | Summary busco_diptera_odb12 |
|---|---|---|---|
| Sylvicola cinctus | Anisopodidae (outgroup) | GCA_963854165.1 | C:88.4%[S:87.5%,D:1.0%],F:4.7%,M:6.9%,n:5067 |
| Bibio marci | Bibionidae | GCA_910594885.2 | C:88.9%[S:88.3%,D:0.6%],F:6.1%,M:5.1%,n:5067 |
| Dilophus febrilis | Bibionidae | GCA_958336335.1 | C:83.1%[S:82.7%,D:0.4%],F:7.1%,M:9.9%,n:5067 |
| Penthetria funebris | Bibionidae | GCA_027564355.1 | C:70.8%[S:70.5%,D:0.3%],F:14.9%,M:14.3%,n:5067 |
| Plecia longiforceps | Bibionidae | GCA_041753815.1 | C:83.7%[S:82.0%,D:1.8%],F:6.3%,M:9.9%,n:5067 |
| Bolitophila cinerea | Bolitophilidae | GCA_010015015.2 | C:82.1%[S:81.4%,D:0.7%],F:9.9%,M:8.0%,n:5067 |
| Bolitophila hybrida | Bolitophilidae | GCA_027564075.1 | C:80.8%[S:80.0%,D:0.8%],F:10.3%,M:8.9%,n:5067 |
| Aphidoletes aphidimyza | Cecidomyiidae | GCA_030463065.1 | C:91.3%[S:89.7%,D:1.6%],F:1.7%,M:7.0%,n:5067 |
| Catotricha subobsoleta | Cecidomyiidae | GCA_011634745.2 | C:84.9%[S:82.8%,D:2.1%],F:7.2%,M:7.9%,n:5067 |
| Contarinia nasturtii | Cecidomyiidae | GCA_009176525.2 | C:89.8%[S:87.7%,D:2.1%],F:2.5%,M:7.6%,n:5067 |
| Lestremia cinerea | Cecidomyiidae | GCA_027564135.1 | C:77.1%[S:22.4%,D:54.7%],F:10.7%,M:12.1%,n:5067 |

| | | | |
|---|---|---|---|
| Mayetiola destructor | Cecidomyiidae | GCA_000149185.1 | C:80.9%[S:79.8%,D:1.1%],F:6.4%,M:12.7%,n:5067 |
| Obolodiplosis robiniae | Cecidomyiidae | GCA_028476595.1 | C:90.6%[S:88.4%,D:2.2%],F:1.8%,M:7.6%,n:5067 |
| Porricondyla nigripennis* | Cecidomyiidae | GCA_026546635.1 | C:39.0%[S:30.3%,D:8.7%],F:21.9%,M:39.1%,n:5067 |
| Resseliella maxima | Cecidomyiidae | GCA_029041755.1 | C:90.8%[S:88.1%,D:2.7%],F:1.9%,M:7.3%,n:5067 |
| Sitodiplosis mosellana | Cecidomyiidae | GCA_021018905.1 | C:84.3%[S:82.8%,D:1.5%],F:2.2%,M:13.6%,n:5067 |
| Diadocidia ferruginosa | Diadocidiidae | idDiaFerr1 | C:92.2%[S:90.1%,D:2.1%],F:3.6%,M:4.2%,n:5067 |
| Symmerus nobilis | Ditomyiidae | GCA_027564815.1 | C:64.2%[S:63.8%,D:0.4%],F:16.8%,M:19.0%,n:5067 |
| Macrocera phalerata | Keroplatidae | GCA_idMacPhal | C:90.5%[S:88.1%,D:2.4%],F:4.0%,M:5.5%,n:5067 |
| Macrocera vittata* | Keroplatidae | GCA_024741295.1 | C:55.0%[S:54.5%,D:0.5%],F:23.0%,M:22.1%,n:5067 |
| Platyura marginata | Keroplatidae | GCA_026745375.1 | C:85.8%[S:85.3%,D:0.6%],F:7.6%,M:6.6%,n:5067 |
| Acnemia nitidicollis | Mycetophilidae | idAcnNiti2 | C:91.6%[S:90.8%,D:0.7%],F:3.9%,M:4.5%,n:5067 |
| Coelosia flava | Mycetophilidae | idCoeFlav1 | C:90.3%[S:89.6%,D:0.7%],F:4.6%,M:5.1%,n:5067 |
| Gnoriste bilineata | Mycetophilidae | GCA_026546565.1 | C:73.7%[S:73.0%,D:0.6%],F:14.1%,M:12.3%,n:5067 |
| Leia winthemii | Mycetophilidae | GCA_965152755.1 | C:88.8%[S:88.0%,D:0.7%],F:5.3%,M:6.0%,n:5067 |
| Bradysia alpicola* | Sciaridae | GCA_049901615.1 | C:46.9%[S:46.2%,D:0.6%],F:18.2%,M:35.0%,n:5067 |
| Bradysia confinis | Sciaridae | GCA_047370885.1 | C:74.1%[S:73.0%,D:1.1%],F:12.2%,M:13.7%,n:5067 |
| **Bradysia coprophila** | Sciaridae | GCA_965233685.1 | C:97.1%[S:54.3%,D:42.7%],F:1.1%,M:1.8%,n:5067 |
| Bradysia desolata* | Sciaridae | GCA_047370855.1 | C:39.8%[S:38.9%,D:0.9%],F:19.6%,M:40.6%,n:5067 |
| **Bradysia impatiens** | Sciaridae | GCA_965233325.1 | C:94.8%[S:72.8%,D:22.0%],F:2.0%,M:3.3%,n:5067 |
| Bradysia nitidicollis | Sciaridae | idBraNiti1 | C:93.5%[S:91.5%,D:2.0%],F:1.9%,M:4.6%,n:5067 |
| Bradysia ocellaris | Sciaridae | idBraOcel1 | C:94.3%[S:92.4%,D:1.9%],F:2.3%,M:3.4%,n:5067 |

| | | | |
|---|---|---|---|
| Bradysia odoriphaga | Sciaridae | GCA_016920775.1 | C:91.9%[S:89.7%,D:2.2%],F:2.6%,M:5.5%,n:5067 |
| Bradysia pectoralis* | Sciaridae | GCA_047370905.1 | C:52.1%[S:51.3%,D:0.7%],F:19.1%,M:28.8%,n:5067 |
| Corynoptera forcipata | Sciaridae | idCorForc1 | C:94.9%[S:91.9%,D:3.0%],F:2.0%,M:3.1%,n:5067 |
| Lycoriella agraria* | Sciaridae | GCA_047370815.1 | C:35.3%[S:35.0%,D:0.3%],F:22.8%,M:41.9%,n:5067 |
| **Lycoriella ingenua** | Sciaridae | GCA_965233655.1 | C:96.1%[S:71.0%,D:25.0%],F:1.7%,M:2.3%,n:5067 |
| Phytosciara flavipes | Sciaridae | idPhyFlap3 | C:94.0%[S:91.2%,D:2.8%],F:2.3%,M:3.7%,n:5067 |
| Phytosciara flavipes | Sciaridae | GCA_026413405.1 | C:60.8%[S:47.5%,D:13.3%],F:19.2%,M:19.9%,n:5067 |
| Pseudolycoriella hygida | Sciaridae | GCA_029228625.1 | C:85.5%[S:82.9%,D:2.5%],F:5.9%,M:8.6%,n:5067 |
| Schwenckfeldina carbonaria | Sciaridae | idSchCarb1 | C:94.6%[S:93.1%,D:1.5%],F:2.2%,M:3.2%,n:5067 |
| Trichosia splendens* | Sciaridae | GCA_026413465.1 | C:46.1%[S:45.3%,D:0.8%],F:25.0%,M:28.9%,n:5067 |

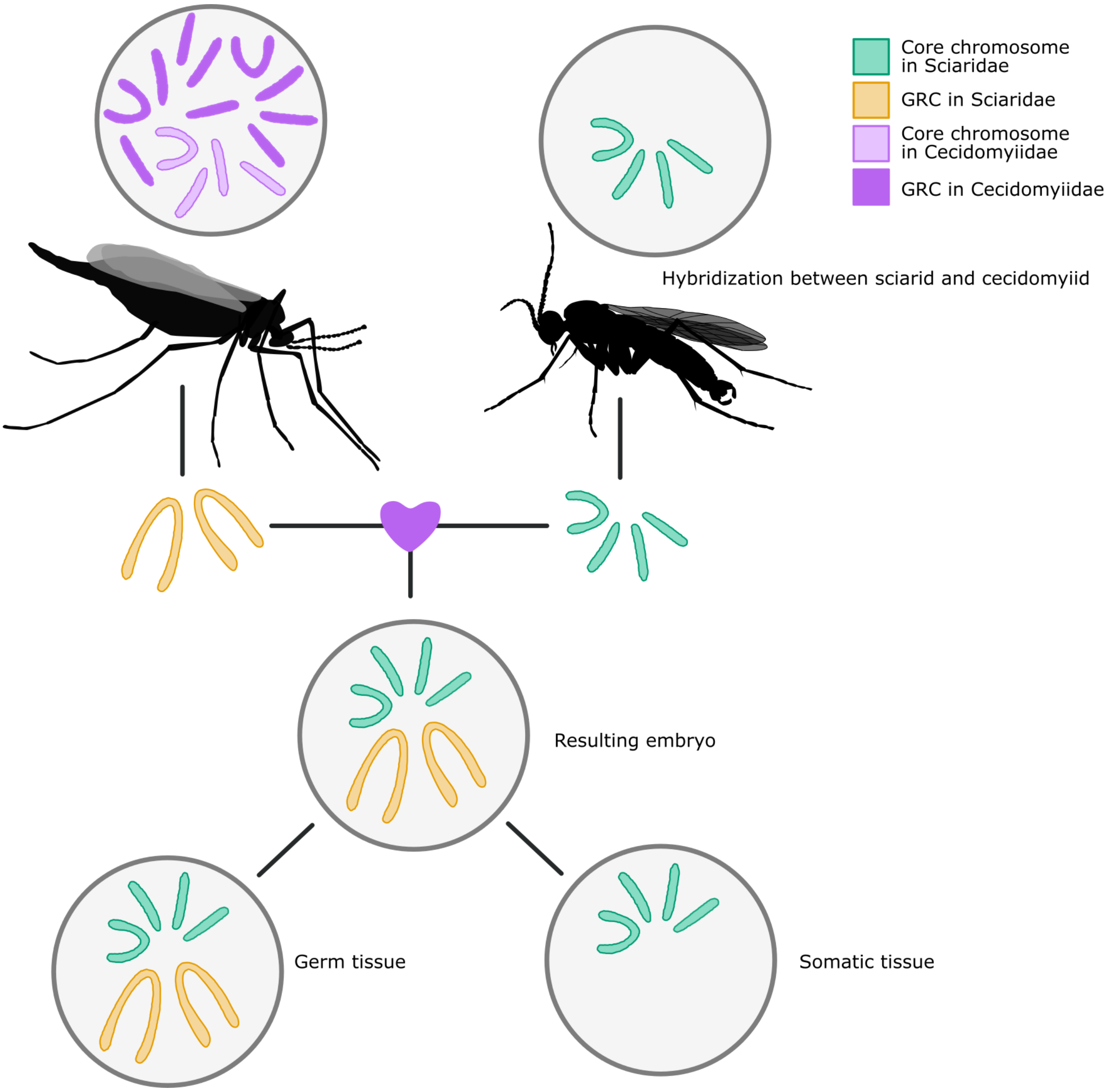


**Supplementary Figure 1.** Scheme of how hybridization between a cecidomyiid and sciarid may have occurred. The sciarid (i.e. the common ancestor of all sciarids) donated the core chromosomes (autosomes/ X chromosome- green) to the embryo which resulted from the hybridization event. Meanwhile, the cecidomyiid ancestor donated the GRCs (coloured orange as we do know whether they originated from cecidomyiid GRCs or core chromosomes) to the embryo. A remaining question is how the core chromosomes retained their diploid state.

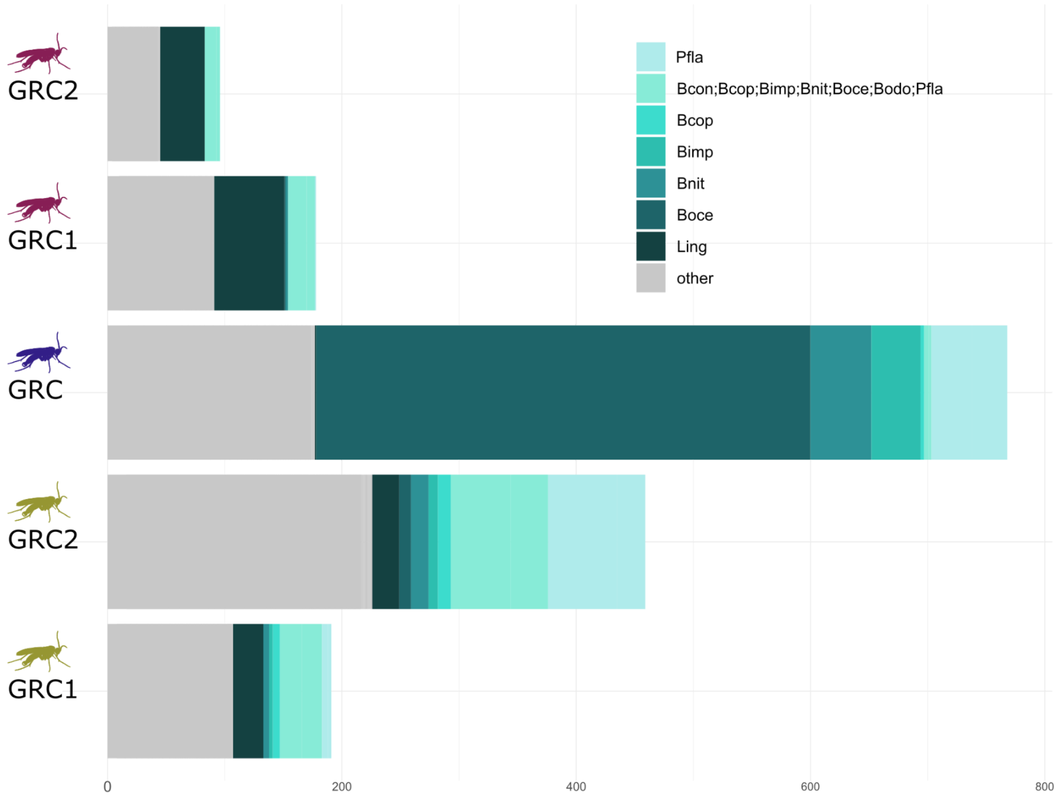


**Supplementary Figure 2.** Nearest non-GRC neighbour in gene trees made from BUSCO genes on GRCs of sciarid origin. For *L. ingenua* GRCs (top two bars), the most frequent non-GRC neighbour is the core gene from *L. ingenua*, for *B. impatiens* (middle), the most frequent non-GRC neighbour is *B. ocellaris* (a closely related species), but for *B. coprophila* (bottom two bars), the most frequent non-GRC neighbour is a clade containing all *Bradysia* species and the two *P. flavipes* genomes (which are combined here are they always formed a clade in gene trees). For each category in the legend, the species involved are designated by the first letter of the genus name followed by the first three letters of the species name.

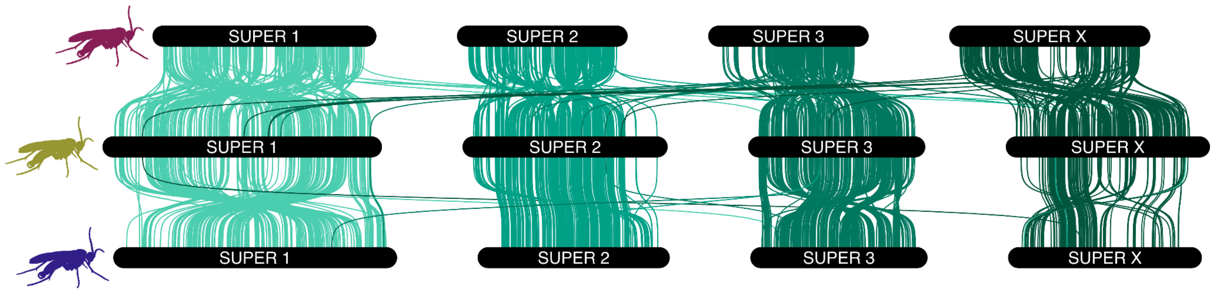


**Supplementary Figure 3.** Synteny between the core chromosomes in *L. ingenua* (top), *B. coprophila* (middle), and *B. impatiens* (bottom). There is clear synteny between the core chromosomes in each species.

### *B. coprophila*

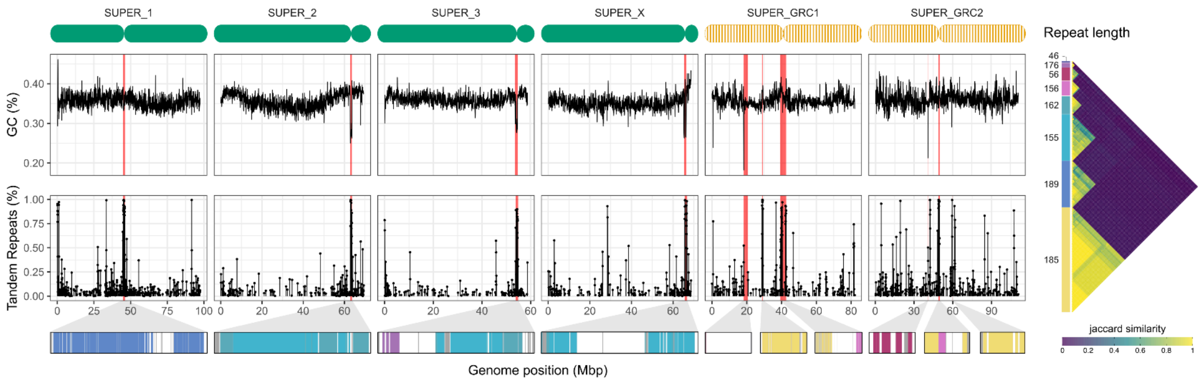


### *B. impatiens*

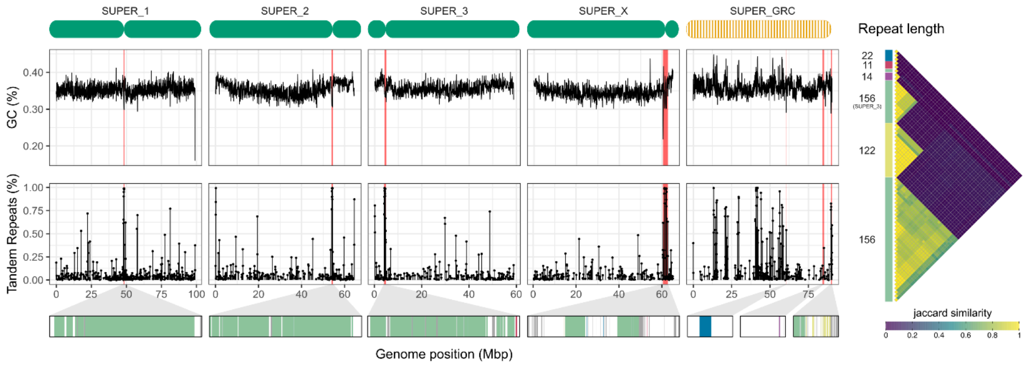


### *L. ingenua*

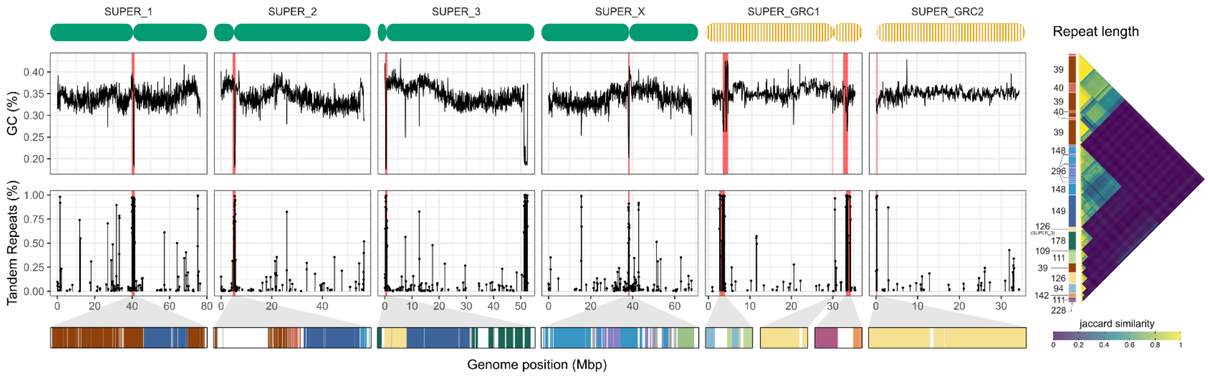


**Supplementary Figure 4**. Centromere annotation. For each species, each chromosome in the genome is shown on the top of the figure with the inferred centromere position indicated. The second panel shows the GC% across the chromosome (in 50-kbp windows) with dips in GC% highlighted red indicating potential centromere position. The next panel shows the tandem repeats density across each chromosome, with regions with a high proportion of tandem repeats highlighted red, indicating possible centromere position. Below that shows an inset of the satellite repeats in the likeliest centromere candidate regions. The right side of the figure shows the monomer repeat size of each of the satellites with a jaccard similarity plot showing sequence similarity between different satellite repeats.

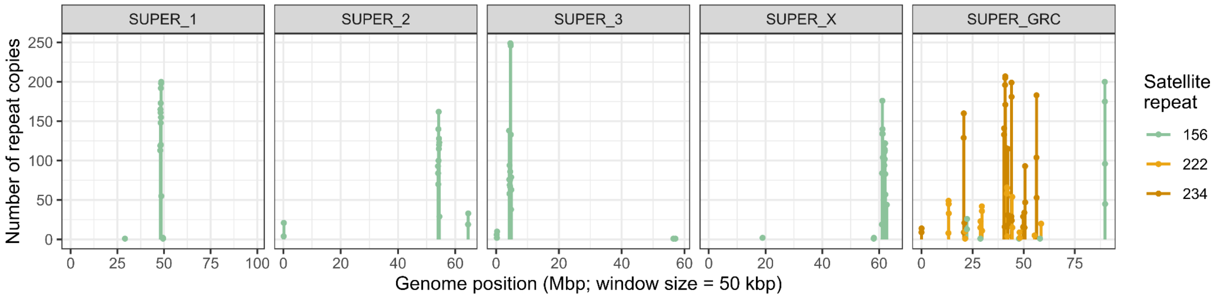


**Supplementary Figure 5**. Main satellite repeats in *B. impatiens*. The GRC in *B. impatiens* has a chromosome-specific 222/234 bp satellite DNA found in high copy number in the middle of the chromosome (probably an old centromere), and in lower copy numbers throughout the rest of the chromosome. We believe the centromere in the GRC in *B. impatiens* is at the end of the chromosome, where the 156 satellite repeat is located

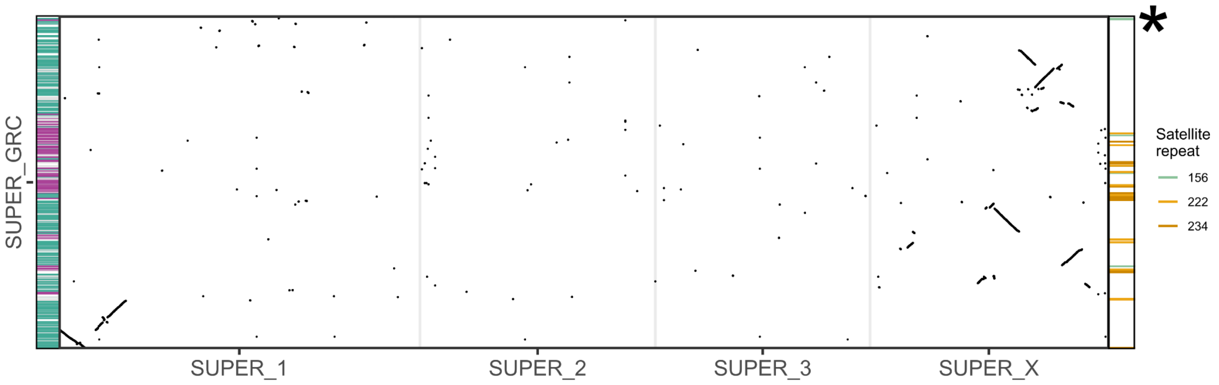


**Supplementary Figure 6.** Synteny between the GRC in *B. impatiens* and the core chromosomes. The centromere is at the end of the GRC (marked with asterisks), right between two regions with different origins. Given that the part of the GRC before that is syntenic to the X chromosome, we believe that the new centromere might have arisen from a duplication of the X chromosome to the GRC, followed by a centromere shift on the GRC.

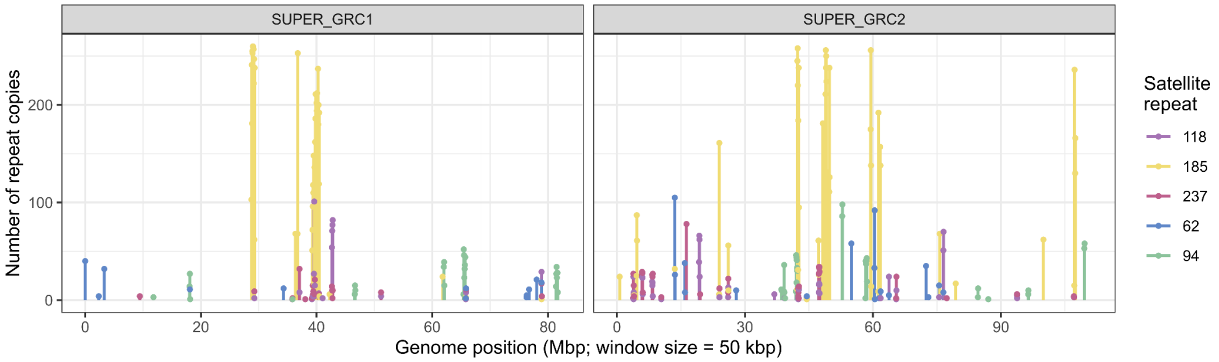


**Supplementary Figure 7.** Positions of the satellite repeats from the short arm of the chromosome X in the GRCs in *B. coprophila*. Unlike in the short X arm, the satellite repeats on the GRCs are found throughout the whole length of the chromosomes.

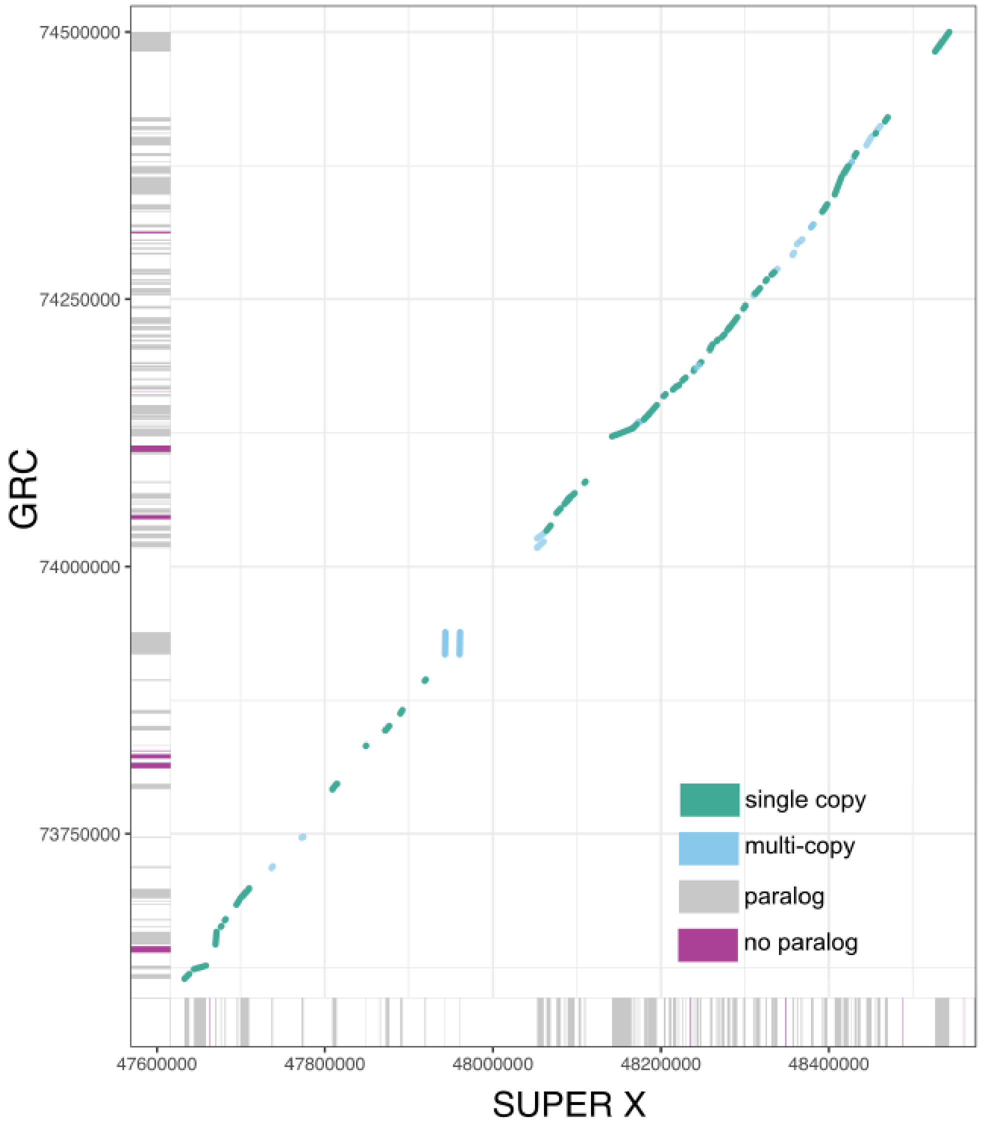


**Supplementary Figure 8.** An example of a syntenic block between the X chromosome and GRC in *B. impatiens*. The majority of genes along the syntenic block are single copy paralogs between the core genome and GRC. Very few genes on the GRC are missing on the syntenic portion of the X chromosome (purple- little evidence of gene degradation on the GRC over time). The conservation of gene order between the X chromosome and GRC is suggestive that this whole block of sequence duplicated from the X chromosome (as genes in this region are in the Sciaridae clade in the phylogenetic analysis) onto the GRC.

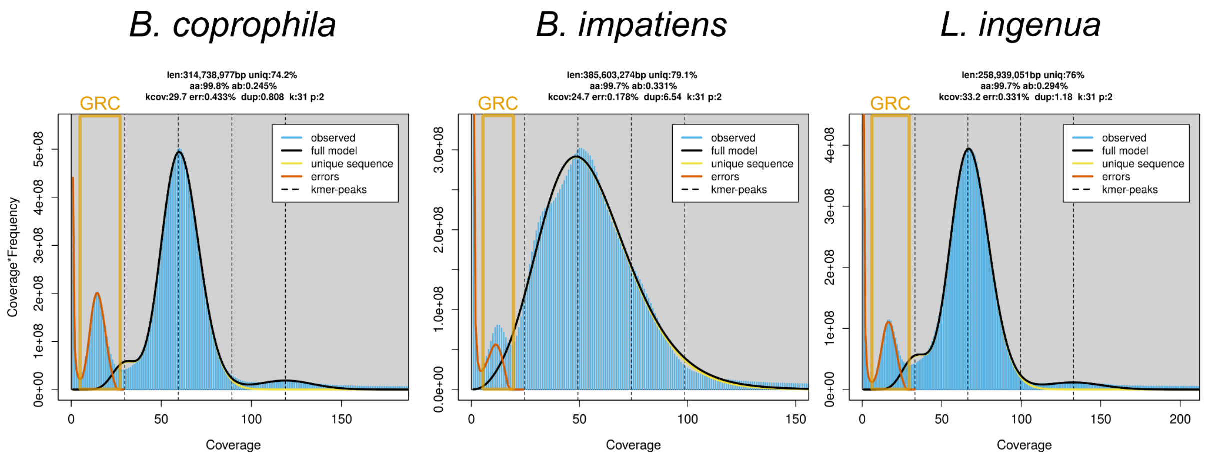


**Supplementary Figure 9.** Genomescope plots for the raw Pacbio data from *B. coprophila*, *B. impatiens* and *L. ingenua*. Because adult male germ tissue is made up of somatic gonad tissue and sperm (germline), GRCs are at a lower frequency in the data compared to other chromosomes (less than half the coverage) and can be easily identified in the genomescope profiles.

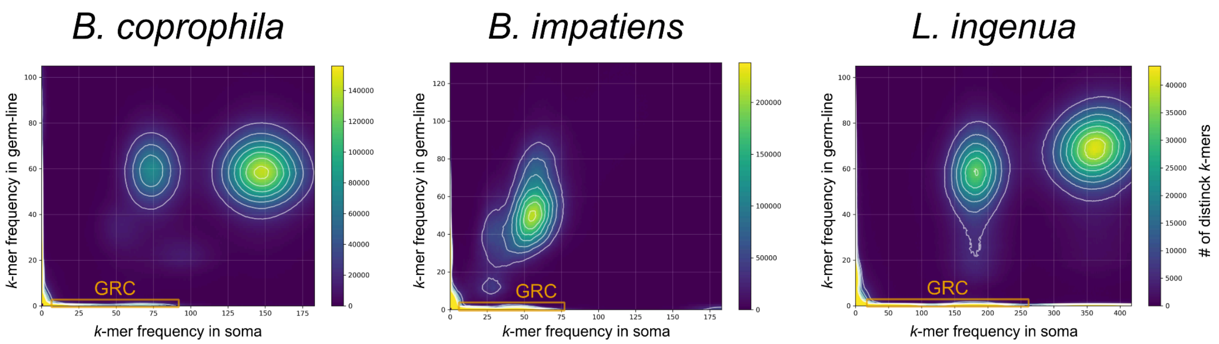


**Supplementary Figure 10.** K-mer comparison plots comparing our Pacbio germline libraries to publically available somatic data. In all cases, you can see the GRC sequence present in the germline library but absent from the somatic library.

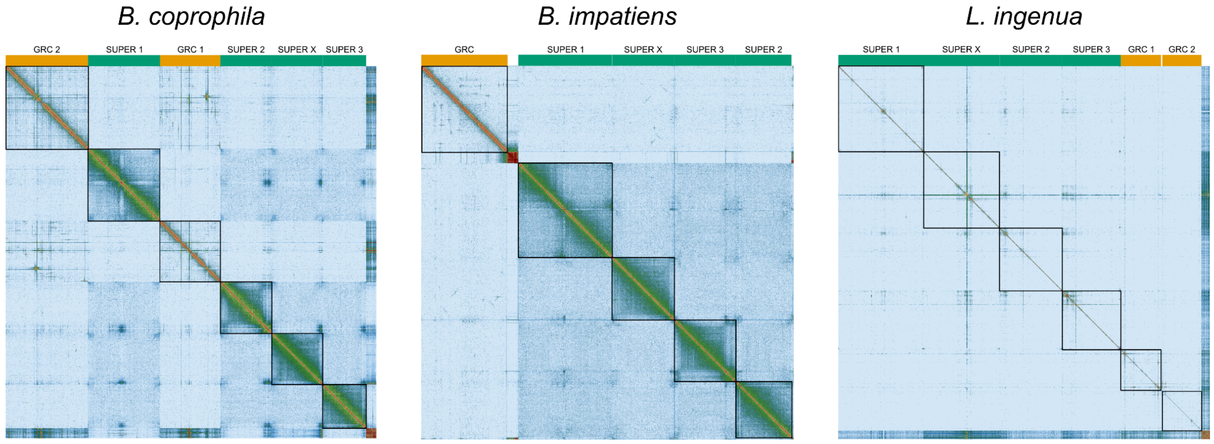


**Supplementary Figure 11.** Hi-C contact maps for *B. coprophila*, *B. impatiens,* and *L. ingenua.* The GRCs are labelled in orange at the top of the map, with the core chromosomes (autosomes and X chromosome) labelled with green. *Bradysia coprophila* has two large GRCs, *B. impatiens* has one large GRC, and *L. ingenua* has two small GRCs.
